# Microbiota metabolized Bile Acids accelerate Gastroesophageal Adenocarcinoma via FXR inhibition

**DOI:** 10.1101/2024.06.11.598405

**Authors:** Theresa Baumeister, Andrea Proaño-Vasco, Amira Metwaly, Karin Kleigrewe, Alexander Kuznetsov, Linus Schömig, Martin Borgmann, Mohammed Khiat, Akanksha Anand, Katrin Böttcher, Dirk Haller, Andreas Dunkel, Veronika Somoza, Sinah Reiter, Chen Meng, Robert Thimme, Roland M. Schmid, Deepa T. Patil, Elke Burgermeister, Yiming Huang, Yiwei Sun, Harris H. Wang, Timothy C. Wang, Julian A. Abrams, Michael Quante

## Abstract

**Background:** The incidence of Barrett esophagus (BE) and Gastroesophageal Adenocarcinoma (GEAC) correlates with obesity and a diet rich in fat. Bile acids (BA) support fat digestion and undergo microbial metabolization in the gut. The farnesoid X receptor (FXR) is an important modulator of the BA homeostasis. The capacity of inhibiting cancer-related processes when activated, make FXR an appealing therapeutic target. In this work, we assess the role of diet on the microbiota-BA axis and evaluate the role of FXR in disease progression.

**Results:** Here we show that high fat diet (HFD) accelerated tumorigenesis in L2-IL1B mice (BE- and GEAC- mouse model) while increasing BA levels and enriching gut microbiota that convert primary to secondary BA. While upregulated in BE, expression of FXR was downregulated in GEAC in mice and humans. In L2-IL1B mice, FXR knockout enhanced the dysplastic phenotype and increased Lgr5 progenitor cell numbers. Treatment of murine organoids and L2-IL1B mice with the FXR agonist obeticholic acid (OCA) deacelerated GEAC progression.

**Conclusion:** We provide a novel concept of GEAC carcinogenesis being accelerated via the diet-microbiome-metabolome axis and FXR inhibition on progenitor cells. Further, FXR activation protected with OCA ameliorated the phenotype in vitro and in vivo, suggesting that FXR agonists have potential as differentiation therapy in GEAC prevention.

**Statement of significance:** If its inhibition is linked to disease progression and its activation to cancer prevention, exploring the potential of FXR as a therapeutic target has great clinical relevance in GEAC context.

## INTRODUCTION

It has been proposed that chronic reflux of gastric and bile acids (BA) triggers inflammation at the gastroesophageal junction (GEJ), leading to Barrett esophagus (BE) and gastroesophagel adenocarcinoma (GEAC) (*1*). High fat, western-style diets and obesity have emerged as risk factors for GEAC (*2, 3*) and represent modifiable targets for cancer prevention. Western-style diet has been linked to the occurrence of precancerous gastrointestinal lesions (*4*), as it can provoke chronic inflammation in the gastrointestinal tract (*5*) and directly affect the composition of gastrointestinal microbiota.

Thus far, studies relating diet and obesity to GEAC generally fail proving direct causality. In our transgenic mouse model of BE and GEAC (L2-IL1B), both, high fat (HFD) and high fructose diet led to a shift in the gut microbiota and accelerated GEAC-carcinogenesis (*6, 7*), suggesting that the intestinal microbiota has an impact on tissues distant to the gut. Gut bacteria are responsible for BA deconjugation and also for the conversion of primary to secondary bile acids (*8*), which can passively diffuse into the blood circulation across the colonic epithelial surface (*9*). Hence, the gut microbiota profile is closely tied to the BA pool composition and secondary BA levels, which are associated with various diseases (*8*). For instance, in L2-IL1B mice, treatment with deoxycholic acid (DCA), a DNA-damaging (*10*) and carcinogenic secondary BA (*11, 12*), induced substantial aggravation of the phenotype (*13*).

Among the BA-activated receptors, the farnesoid X receptor (FXR) is the most important one as it is the main modulator of BA homeostasis and enterohepatic circulation. Although FXR can bind to many BAs, the downstream effects of FXR vary across BAs. FXR is a nuclear “orphan class” receptor, the main BA receptor and it is primarily expressed in the small intestine and the liver. Upon ligand-mediated activation, FXR forms a heterodimer with the retinoid-x-receptor (RXR), binds to specific DNA elements and regulates the expression of downstream target genes (*14*). FXR activation is important in BA metabolism and homeostasis. Additionally, if activated, FXR was shown to inhibit intestinal tumorigenesis (*15*), hepatic tumor cell proliferation, dedifferentiation and migration (*16*). It has also been described to inhibit inflammatory signaling (*17*). Moreover, in a colorectal cancer model, FXR expression was downregulated upon HFD exposure and treatment with an FXR agonist ameliorated the phenotype of these mice (*18*).

The manifold functions of FXR intervening in metabolic and tumorigenic signaling make FXR an attractive therapeutic target for treatment of BA-mediated metabolic and gastrointestinal diseases (*19*). In addition, as a first drug targeting FXR, obeticholic acid (OCA), a semisynthetic, highly specific FXR agonist was approved for treatment of primary biliary cholangitis. In light of the continued rise in GEAC incidence, safe and effective chemopreventive agents could have a significant public health impact.

Here, we analyze the effects of dietary-modulated intestinal microbiota on GEAC carcinogenesis utilizing the L2-IL1B mouse model and assessed the BA signatures of healthy controls, BE- and GEAC-patients. We provide evidence of diet-related changes in the intestinal microbiota composition having an impact on BA metabolism and FXR levels and hence, on cancer-associated pathways in the metaplastic BE regions in mice and humans. Our results outline a novel concept of how the diet-microbiota-bile acid axis can affect tumorigenesis.

## MATERIALS AND METHODS

### Transgenic mouse strains

All animal experimental work performed in Germany was carried out under the approval of the district government of Upper Bavaria, according to the animal experimental approvals 55.2.1.54-2532-125-12 and 55.2-1-54-2532-24-2016. All animal experiments were conducted in accordance with Animal Welfare and Ethical Guidelines of the Klinikum Rechts der Isar, TUM, Munich and the Columbia University, New York. All animal experimental work in this study was based on the transgenic mouse model (L2-IL1B) of BE and GEAC (*20*). L2-IL1B mice overexpress the human interleukin 1 beta in the esophageal and squamous forestomach mucosa under control of the Epstein-Barr-Virus promoter (EBV-L2). L2-IL1B mice develop esophagitis with subsequent progression to BE and ultimately to GEAC without any additional intervention. To obtain L2-IL1B-FXR KO mice, L2-IL1B mice were backcrossed to C57BL/6J mice and crossed with FXR KO mice, which have a loxP-Cre-based whole-body knockout of FXR(*21*).

### Dietary treatment of L2-IL1B mice

For dietary treatment studies in Germany, starting at 8-9 weeks of age, mice were randomly distributed in treatment groups and fed high fat diet (HFD, Ssniff, S5745-E712), a matching control diet (CD, Ssniff, S5745-E702) or high fat diet / control diet + obeticholic acid (HFD+OCA, S0615-E710; CD+OCA, Ssniff, S0615-E705). For dietary treatment studies in New York, starting at 8-9 weeks of age, mice were randomly distributed in treatment groups and fed CD, HFD or HFD+OCA (dietary composition analogous to the diets used in Germany).

### Human BarrettNET study cohort

Samples and data from a prospective study of 49 patients with non-dysplastic BE (BE), low grade dysplasia (LGD), high grade dysplasia (HGD), or GEAC, as well as non-BE controls; a subset of patients from the BarrettNET study at Klinikum Rechts der Isar were analyzed for changes in bile acid metabolizing bacteria or feces and serum bile acids with disease progression(*22*).

### Patient and Public Involvement

Patients were asked to evaluate the process of BarrettNET within a QM questionnaire and individual study results were communicated to the patient and used for diagnostic and therapeutic evaluation.

### Statistical analysis

Statistical analyses were performed using GraphPad Prism version 8.00 for Windows (GraphPad Software). Detailed description of the tests used during statistical analysis to be found in the supplementary methods.

**Further materials and methods can be found in the Supplements.**

## RESULTS

### By shifting the composition of the gut microbiota and the resulting BA profile, HFD facilitates the exposure of BE tissue to harmful BA

We previously demonstrated that HFD induced alterations to the gut microbiota and accelerated the dysplastic phenotype in L2-IL1B mice (*6*). Here, we found that mice fed HFD had decreased relative abundance of Bacteroidota, a phyla containing taxa with BA-deconjugating capacities (*23*), and a significant increase in bacteria with BA-converting capabilities (*23-25*) (Fig.1 A). Moreover, diet shifted the concentration of certain BAs in gut tissue, feces, serum and BE tissue (Fig. 1B-E). Lithocholic acid (LCA) and Chenodeoxycholic acid (CDCA) have been described to contribute to carcinogenesis in the esophagus (*26*). HFD increased levels of their conjugates 12-Keto-LCA and Tauro-CDCA in feces, suggesting a potential carcinogenic effect if translocated from the gut lumen into the esophagus. Moreover, in BE tissue of the mice fed HFD, levels of Isodeoxycholic acid (Iso-DCA) and Taurocholic acid (TCA) were significantly increased. While TCA has been described to promote growth in vitro, DCA has been linked to esophageal carcinogenesis by inducing pathways of inflammation and DNA damage (*27*). Taken together, these data support the concept that HFD promotes GEAC in part, by changing the BA pool in the gut. We hypothesize that BA can be then absorbed from the gut and reach distal tissue through blood circulation.

**Fig. 1.**
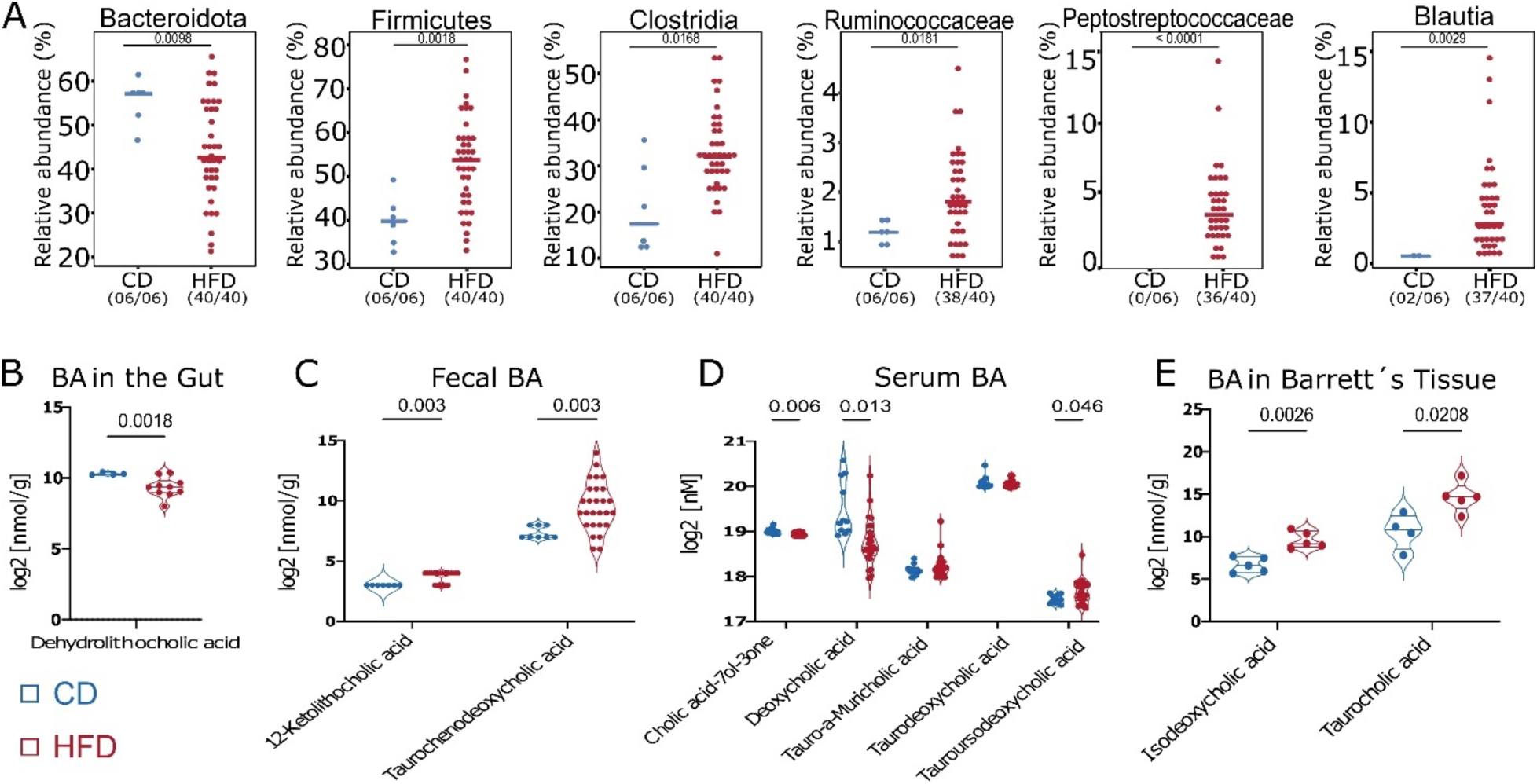
HFD alters BA metabolism by enrichment of bacteria with BA-converting capabilities leading to a shift in the BA profile in feces, serum, local and distal tissue. **(A)** Relative abundance of bacteria with BA-deconjugating and BA-converting capabilities differs between CD vs. HFD. If significantly different, adjusted p-value is shown. **(B)** Log_2_ transformed concentration of Dehydrolithocholic acid in intestinal tissue of mice fed HFD (n=10) decreased in comparison to mice fed CD (n=4). **(C)** Log_2_ transformed concentration of 12-Ketolithocholic acid and Taurochenodeoxycholic acid in feces of mice fed HFD (n=27) increased in comparison to mice fed CD (n=8). **(D)** Log_2_ transformed concentration of single BA in sera of mice fed HFD and CD. While Cholic acid-7ol-3-one and Deoxycholic acid decreased mice fed HFD, Tauro-alpha-Muricholic acid, Taurodeoxycholic acid and Tauroursodeoxycholic acid increased in mice fed HFD (n=24) compared to mice fed CD (n=11). **(E)** Log_2_ transformed concentration of Isodeoxycholic acid and Taurocholic acid in Barrett’s tissue of mice fed HFD (n=5) was increased in comparison to mice fed CD (n=5). For A, fecal samples of 3-12 month-old L2-IL1B mice were combined. Data are presented as mean with SD. For B-E, samples of 6-9-month-old L2-IL1B mice were combined.

### Loss of FXR aggravates dysplasia

An important modulator in BA metabolism is FXR. Analysis of previous mRNA expression data of human and mouse (*28*) demonstrate that FXR is upregulated in BE but not in GEAC (Fig. 2A). This was confirmed by immunofluorescence of human BE and GEAC organoids when stained for FXR (Fig. 2B-C) and by immunohistochemstry in the GEJ of L2-IL-1B mice (Fig. 2D). Compared to L2-IL1B mice, L2-IL1B mice with whole body FXR knockout (L2-IL1B-FXR KO) had increased inflammation, dysplasia and tumor formation with associated decreased differentiation reflected by the reduction of goblet cells (Fig. 2E-H). Abrogation of FXR signaling correlated with reduced PAS-positive mucus-producing cells, increased caspase 1 inflammasome activation, α-SMA positive fibroblast infiltration, DNA damage and increased numbers of Lgr5 progenitor cellsat the GEJ (Fig. 2I-O). Importantly, we observed co-localization of Lgr5 and FXR in columnar cells at the GEJ in L2-IL1B mice and an upregulation of stem cell signaling pathways upon loss of FXR, pointing to a direct impact of FXR on Lgr5+ progenitor cells (Fig. 2M-P). Transcriptome comparisons between L2-IL1B and L2-IL1B-FXR KO mice suggested that loss of FXR promoted tissue re-organization and pro-tumorigenic signaling pathways, such as K-RAS and P53 (Supplementary Fig. 1A-C). These data suggest that FXR has a protective effect in BE tissue, and loss of its expression, specifically on progenitor cells, correlates with de-differentiation and disease progression to dysplasia.

**Fig. 2:**
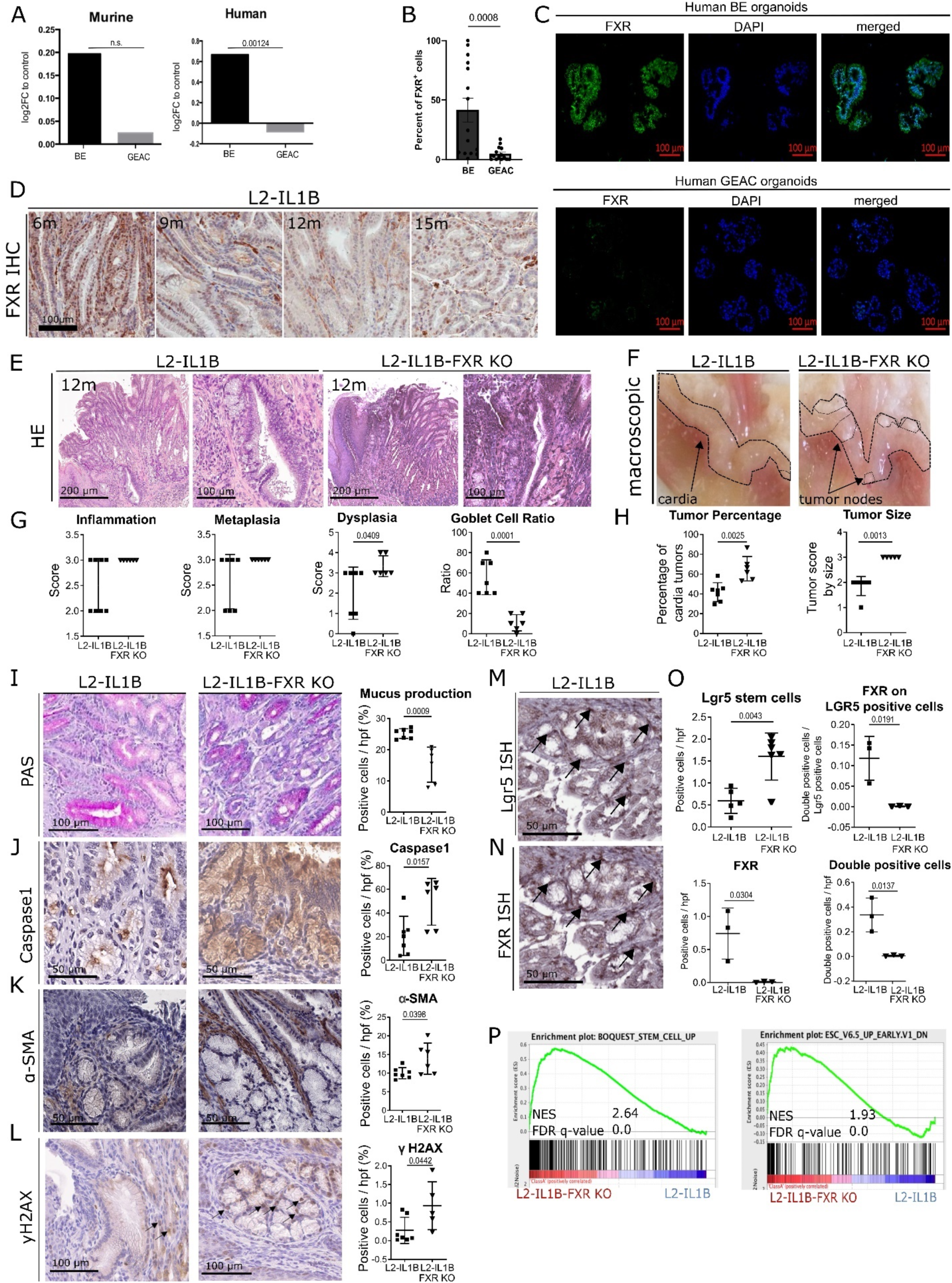
FXR has a protective role in the malignant progression of BE to GEAC. **(A)** Murine FXR expression data of RNA microarray (GSE24931) in adult BE mice and GEAC mice. Compared to normal tissue, expression of FXR in BE-mice was higher and in GEAC lower. Analysis of publicly available gene expression data (*28*) for FXR in human normal, BE or GEAC tissue samples (GSE13898). Expression of FXR in human BE tissue is significantly higher compared to GEAC tissue (adjusted p-value). **(B)** Percentage of FXR positive cells in FFPE-human organoids of BE and GEAC patients. Number of FXR positive cells in BE organoids is significantly higher than in GEAC organoids **(C)** Representative images of FXR immunofluorescence staining of human BE and GEAC organoids **(D)** Representative images of FXR immunohistochemistry staining at the GEJ of L2-IL1B mice (6-, 9-, 12- and 15-months). **(E)** Representative images of HE staining at the GEJ of 12-month-L2-IL1B and L2-IL1B-FXR KO mice **(F)** Representative macroscopic images of the GEJ region of 12-month-L2-IL1B and L2-IL1B-FXR KO mice **(G)** Disease scores as well as goblet cell (GC) ratios of 12-month L2-IL1B (n=7) and L2-IL1B-FXR KO (n=6) mice. While the dysplasia score of L2-IL1B-FXR KO mice was significantly higher than in L2-IL1B mice, the GC ratio was significantly lower. **(H)** Tumor percentage and tumor size were both significantly higher in 12-month L2-IL1B (n=7) compared to L2-IL1B-FXR KO (n=5) mice. **(I)** Representative images of Periodic Acid-Schiff (PAS) staining **(J)** Representative images of Caspase1 immunohistochemistry staining. **(K)** Representative images of alpha-smooth muscle actin (α-SMA) immunohistochemistry staining **(L)** Representative images of yH2AX immunohistochemistry staining **(M)** Lgr5-In-situ-hybridization at the GEJ of 12-month-old L2-IL1B mice **(N)** FXR- In-situ-hybridization at the GEJ of 12-month-old L2-IL1B mice **(O)** Lgr5 positive cells in 12-month L2-IL1B-FXR KO (n=6) compared to L2-IL1B (n=5). FXR positive cells in 12-month L2-IL1B (n=3) compared to L2-IL1B-FXR KO (n=3). Both, FXR expression on LGR5 stem cells and FXR-Lgr5 double positive cells per high power field (hpf) showed higher numbers of double positive cells in L2-IL1B compared to L2-IL1B-FXR KO mouse cardia regions. **(P)** Representative gene sets for a stem-cell like phenotype were enriched in a microarray analysis of 12-month L2-IL1B-FXR KO compared to L2-IL1B mice (gene set enrichment analysis of cardia tissue microarray; n=3). Data are presented as mean with SD or, for metabolomic analysis, mean only. (G-J) Data are displayed as positive cells / hpf= area BE region (μM^2^)/1000 (GEJ of 12-month old mice: L2-IL1B n=7 and L2-IL1B FXR KO n=6). (O) Data are displayed as positive cells / high power field (hpf) = area BE region (μM^2^)/1000 or as FXR-Lgr5 double positive cells per Lgr5 positive stem cells. (12-m-old L2-IL1B n=3-6; 12-m-old L2-IL1B FXR KO n=3-6). (P) Gene expression data was extracted from microarray analysis, and gene set enrichment analysis (GSEA) was performed using the Broad Institute GSEA software.

### Treatment with FXR agonist obeticholic acid (OCA) ameliorates phenotype in vitro

To better understand the effects of FXR, we treated BE organoids isolated from L2-IL1B mice with the secondary and toxic BAs DCA and TβMCA. We combined this treatment with or without the selective FXR agonist OCA. DCA treatment led to cellular and oxidative DNA damage and to lower organoid numbers despite increased proliferation (Fig. 3A-I). TβMCA treatment did not increased DNA damage but increased proliferation (Fig. 3A-D, J-N). As suspected, OCA prevented the toxic and proliferative effects of both secondary bile acids (Fig. 3A-D, G, K-L). To examine the effect of OCA on disease progression in vivo, we treated HFD-fed mice with OCA and compared them to matched CD- and HFD-fed mice. In HFD-fed mice, OCA treatment significantly reduced proliferation, abrogated dysplasia and induced goblet cell (GC) differentiation (Fig. 4A-F). HFD also induced an increase in Lgr5+ and Lgr5+/FXR+ cells at the GEJ (Fig. 4G, Supplementary Fig. 2A). HFD-fed mice treated with OCA treatment also led to reduced levels of neutrophils in BE tissue (Supplementary Fig. 3A-B) and elevated numbers of NK and activated NKT cells (Supplementary Fig. 3C-F). Consequently, we showed *in vitro* and *in vivo*, that the FXR agonist OCA can prevent the acceleration of dysplasia likely due to its effects on the progenitor cell population at the GEJ.

**Fig. 3:**
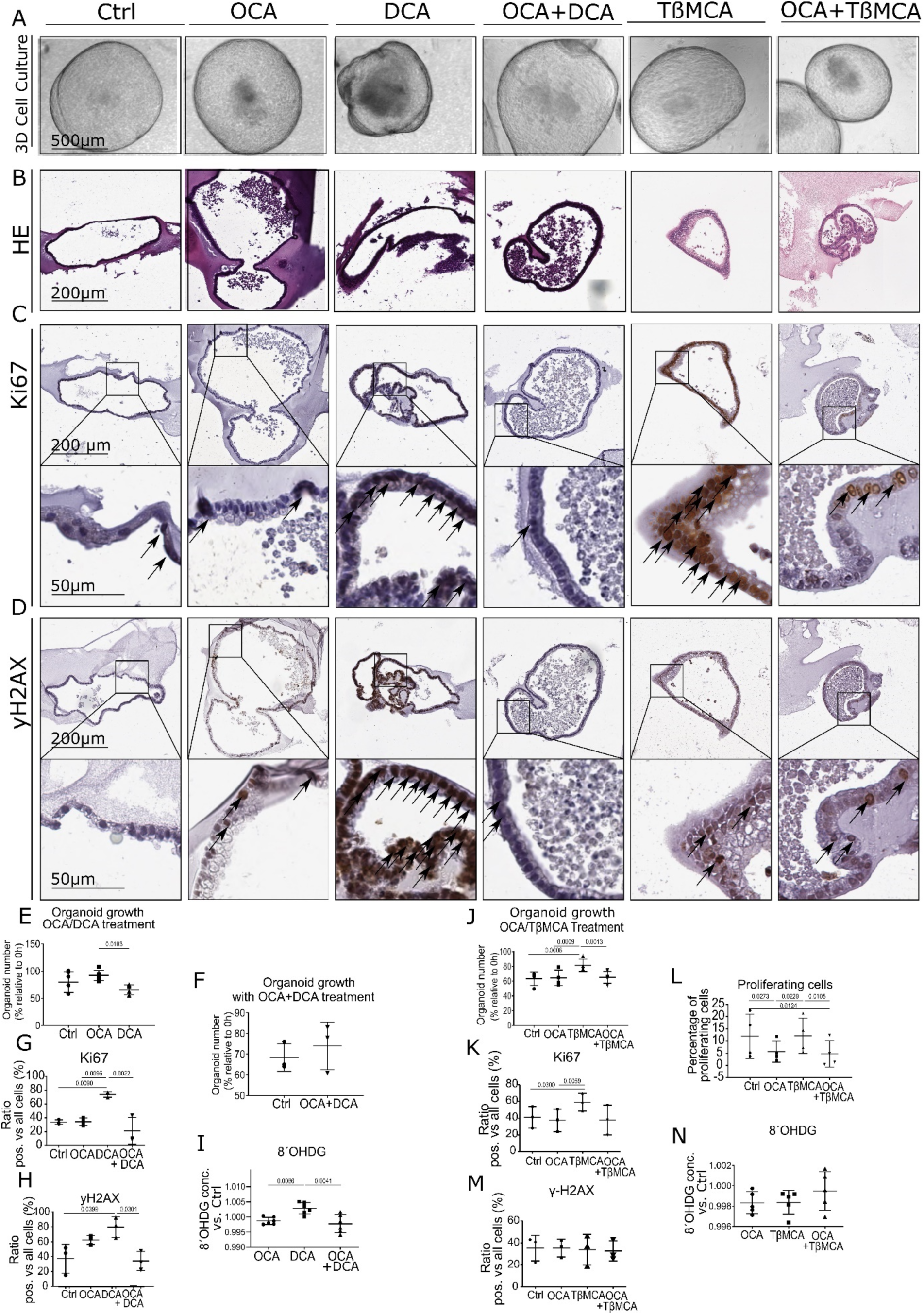
FXR agonist OCA rescues the damaging and proliferative effects of BAs on organoids of L2-IL1B mice. Organoids were treated with 10 µM OCA, DCA, OCA+DCA, TßMCA or OCA+TßMCA in the media for 72 h, respectively. Organoids were counted at 0h and 72h. **(A)** Representative images organoid cultures. Dead cells accumulated in the center of the organoids treated with DCA, and organoids acquired irregular morphologies, whereas organoids treated with OCA or OCA+DCA grew similar to untreated control organoids in terms of size and shape. **(B)** Representative images of HE staining showing the organoid histology **(C)** Representative images of Ki67 immunohistochemistry on organoids **(D)** Representative images of yH2AX immunohistochemistry on organoids **(E)** Quantification of organoid numbers during treatment (10µM OCA, 10µM DCA, untreated as control). While OCA- and DCA-treated samples did not differ from untreated controls, organoid numbers in the samples treated with DCA were lower compared to the samples treated with OCA. **(F)** Quantification of organoid numbers during treatment (10µM OCA + 10µM DCA) compared to untreated organoids. The number of organoids treated with OCA and DCA did not differ compared to controls. **(G)** Ki67 immunohistochemical analysis of organoids showed a higher proliferation in DCA treated samples compared to controls, OCA, and OCA+DCA treated samples (n=3). **(H)** yH2AX immunohistochemical analysis of organoids showed a higher DNA damage in DCA-treated samples compared to controls and OCA+DCA treated samples (n=3). **(I)** 8’OHDG ELISA to determine cellular damage in OCA-, DCA- and OCA+DCA-treated samples normalized to controls showed increased damage in conditioned media of DCA-treated organoids compared to both OCA and OCA+DCA (n=6). **(J)** Number of organoids, depicted as percentage relative to timepoint 0 (0h) and after treatment with 10µM OCA, 10µM TßMCA or 10µM OCA + 10µM TßMCA for 72 h. **(K)** Ki67 immunohistochemical analysis of organoids treated with 10µM of OCA, 10µM TßMCA, 10µM OCA+TßMCA or untreated control. **(L)** Proliferating cells measured with the Click IT FC flow cytometry Kit in organoids after treatment with 10µM OCA, 10µM TßMCA or 10µM OCA and 10µM TßMCA for 72 h. **(M)** yH2AX immunohistochemical analysis of organoids treated with 10µM OCA, 10µM TßMCA and 10µMOCA+TßMCA or untreated controls. **(N)** 8’OHDG ELISA to determine cellular damage in with 10µM OCA-, TßMCA- and OCA+TßMCA-treated samples normalized to controls showed no differences of 8’OHDG in conditioned media between treatments (n=5). Data are presented as mean with SD. (E, F, J) Data are presented as organoid numbers related to 0h timepoint. (G, H, K, M). Data are presented as ratio of positive vs total cells. For (I, N) Data show the concentration of 8’OHDG in conditioned media normalized to control.

**Fig. 4:**
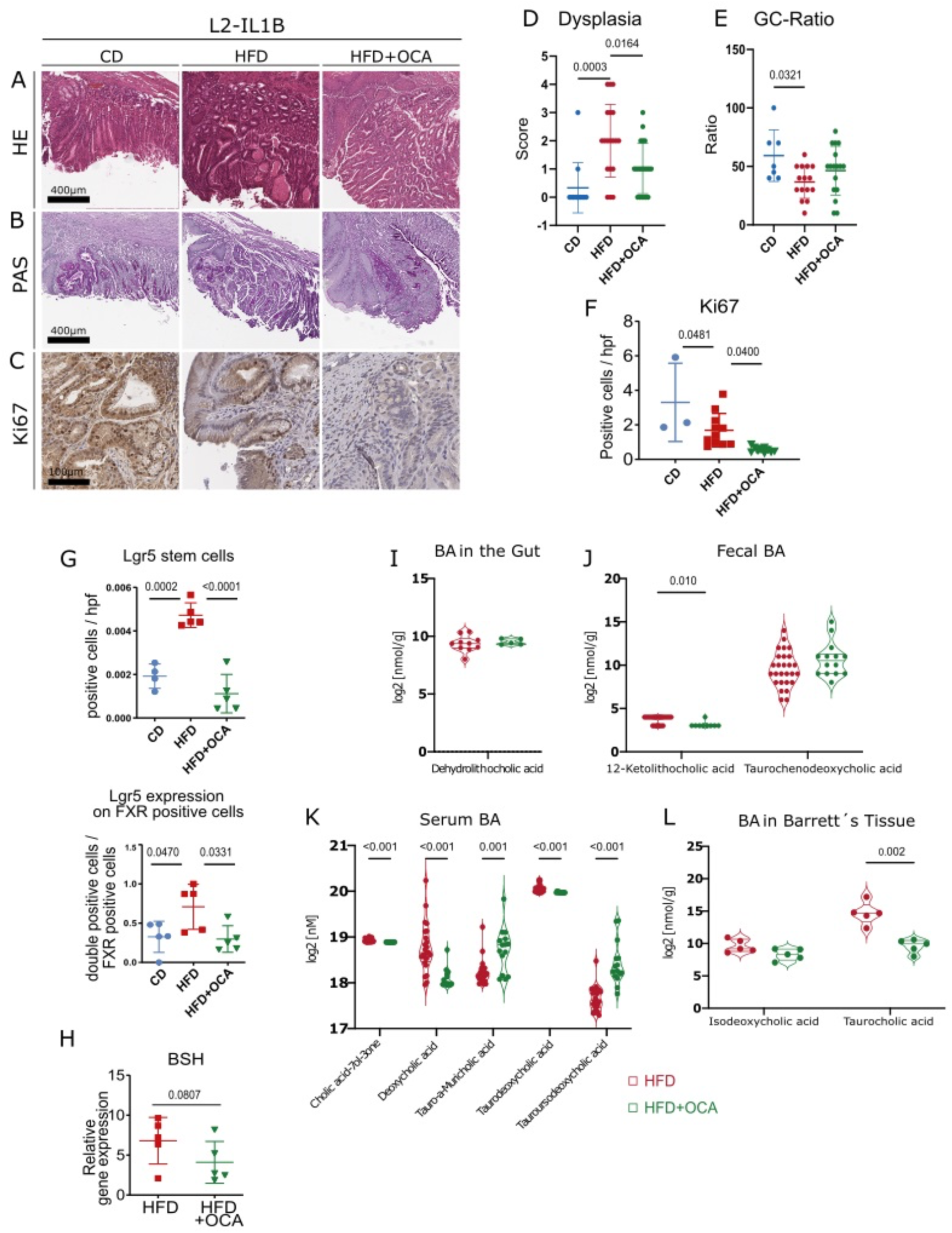
OCA ameliorates the dysplastic phenotype of HFD-fed L2-IL-1B mice and alters the BA profile in feces, serum, local and tissue. **(A)** Representative images of HE staining **(B)** Representative images of PAS staining **(C)** Representative images of Ki67 staining **(D)** Dysplasia scores of L2-IL1B mice (6-9-month old) fed CD (n=12), HFD (n=19) or HFD+OCA (n=20) **(E)** GC ratio of of L2-IL1B mice (6-9-month old) fed CD (n=7), HFD (n=15) or HFD+OCA (n=17) **(F)** Ki67 immunohistochemistry for evaluation of proliferation showed significantly decreasing proliferation rates from CD to HFD and to HFD+OCA (n=4-12). **(G)** Lgr5 positive cells and Lgr5 expression on FXR positive cells in HFD is significantly increased in HFD-fed mice compared to CD and HFD+OCA (n=4-5). **(H)** BSH gene expression tended to be lower in 6-monthL2-IL1B mice fed HFD+OCA compared to mice fed HFD (n=5). **(I)** Log_2_ transformed concentration of Dehydrolithocholic acid in intestinal tissue of mice fed HFD (n=10) compared to mice fed HFD+OCA (n=5) **(J)** Log_2_ transformed concentration of 12-Ketolithocholic acid and Taurochenodeoxycholic acid in feces of mice fed HFD (n=27) compared to mice fed HFD+OCA (n=14) **(K)** Log_2_ transformed concentration of single BA in sera of mice fed HFD and CD. While Cholic acid-7ol-3-one, Deoxycholic acid and Taurodeoxycholic acid decreased in mice fed HFD+OCA, Tauro-alpha-Muricholic acid and Tauroursodeoxycholic acid increased in mice fed HFD+OCA (n=15) compared to mice fed HFD (n=24) **(L)** Log_2_ transformed concentration of Isodeoxycholic acid and Taurocholic acid in Barrett’s tissue of mice fed HFD+OCA (n=5) was increased in comparison to mice fed HFD (n=5) (A-C) Representative images of stainings at the GEJ of L2-IL1B mice fed CD, HFD and HFD+OCA. (F-G) Data are displayed as positive cells / high power field (hpf) = area GEJ (μm^2^)/1000.

### In the L2-IL1B mouse model, OCA decreases dysplasia and proliferation rate at the GEJ while reducing levels of growth-promoting secondary BAs

Considering that metabolic changes in the gut seem to be linked to systemic metabolic changes, we further delved into the microbiota-metabolome-phenotype axis. Bile Salt Hydrolase (BSH) is a microbial, BA-deconjugating enzyme that deconjugates primary BAs, which unconjugated, can be then transformed into secondary BAs and be further re-conjugated (*29*). Expression of BSH was reduced by OCA treatment in HFD-fed mice (Fig. 4H). The fecal bacterial profile of especially of 6-month L2-IL1B mice, changed significantly between HFD-fed and HFD+OCA-fed mice (Supplementary Fig. 4A-B). Microbial analyses showed that bacterial community richness dropped significantly in the feces from 6- to 9-month-HFD-fed mice (Supplementary Fig. 5A).

Additionally, depending on the diet and the age of the mice, a bacterial shift was observed (Supplementary Fig. 5B-J). The cecal microbial profile closely resembled the fecal microbial profile (Supplementary Fig. 4C-D; Supplementary Fig. 6A-I). In both, feces and cecal contents, bacteria belonging to Clostridia, Lachnospiraceae and Ruminicoccaaceae correlated with inflammation, dysplasia and with the BA Taurolithocholic acid, TLCA (Supplementary Fig. 4E, 5L, 6J). TLCA has been described to promote cell growth in Cholangiocarcinoma (*30*) and to promote epithelial-mesenchymal transition (*31*).

A balanced ratio of Firmicutes/Bacteroidetes (F/B) is widely associated with normal intestinal homeostasis and alterations in this ratio are linked to gut dysbiosis and certain diseases (*32*). Here, while Bacteroidetes positively correlated with dysplasia and negatively with GC-ratio, Firmicutes negatively correlated with dysplasia and positively with GC-ratio (Supplementary Fig. 5L).

Moreover, OCA treatment altered the concentration of certain BAs in gut tissue, feces and serum of L2-IL1B mice fed with HFD (Fig. 4I-K). Specifically, OCA treatment decreased serum levels of DCA and Taurodeoxycholic acid, TDCA (Fig. 4K), which have been described to promote progression of GEAC (*27*). OCA treatment also lowered Taurocholic acid (TCA) in BE tissue (Fig. 4L), which has been linked to invasive growth *in vitro (33)*.

Besides linking TLCA and certain bacteria to inflammation and disease, our data suggests that OCA is capable of reducing contents of growth-promoting secondary BAs such as DCA and TDCA, presumably by decreasing the expression of BSH.

### Stool and serum BA levels in BE and GEAC patients correlate with disease stage

To explore the microbiota-metabolome axis at a more clinical level, we analyzed a cohort of patients at different states of esophageal carcinogenesis (non-BE controls, BE, Dysplasia, GEAC) from the BarrettNET Registry (*34*). Bile acid profiling of stool showed clustering linked to disease stage, and enriched conjugated BAs in dysplastic and GEAC-patients (Fig. 5A-B). Despite unclear clustering, bile acid profiling of sera also showed conjugated BAs being enriched in dysplastic and GEAC-patients (Fig. 5C-D). Furthermore, TDCA, Taurochenodeoxycholic acid (TCDCA) and Glycocholic acid (GCA) were enriched in both stool and serum of Dysplastia/GEAC patients. While DCA appears to promote carcinogenesis by inducing pathways of inflammation and DNA damage, CDCA is linked to carcinogenesis by enhancing cell proliferation and promoting of angiogenesis (*27*). Moreover, we identified the bacteria Bifidobacteria, Gremmiger and Ruminococcus to to strongly correlate with the BAs enriched in the stool of Dysplasia/GEAC-patients (Supplementary Fig. 7A). Here we show the expression of BSH to positively correlate with disease severity (Fig. 5E). In the colon, gut bacteria convert the primary BA cholic acid (CA) into the secondary BA DCA (*35*). Interestingly, in patientśsera, levels of these BAs positively correlated with *Lactobacillaceae* (Fig. 5F), and members of this family have been described to express BSH (*36*). Matching with the hypothesis that microbial metabolites in the gut reach distant tissues (*37*) after being absorbed in the gut and entering the circulation, our results show that metabolic changes in the gut can be reflected in the serum.

**Fig. 5:**
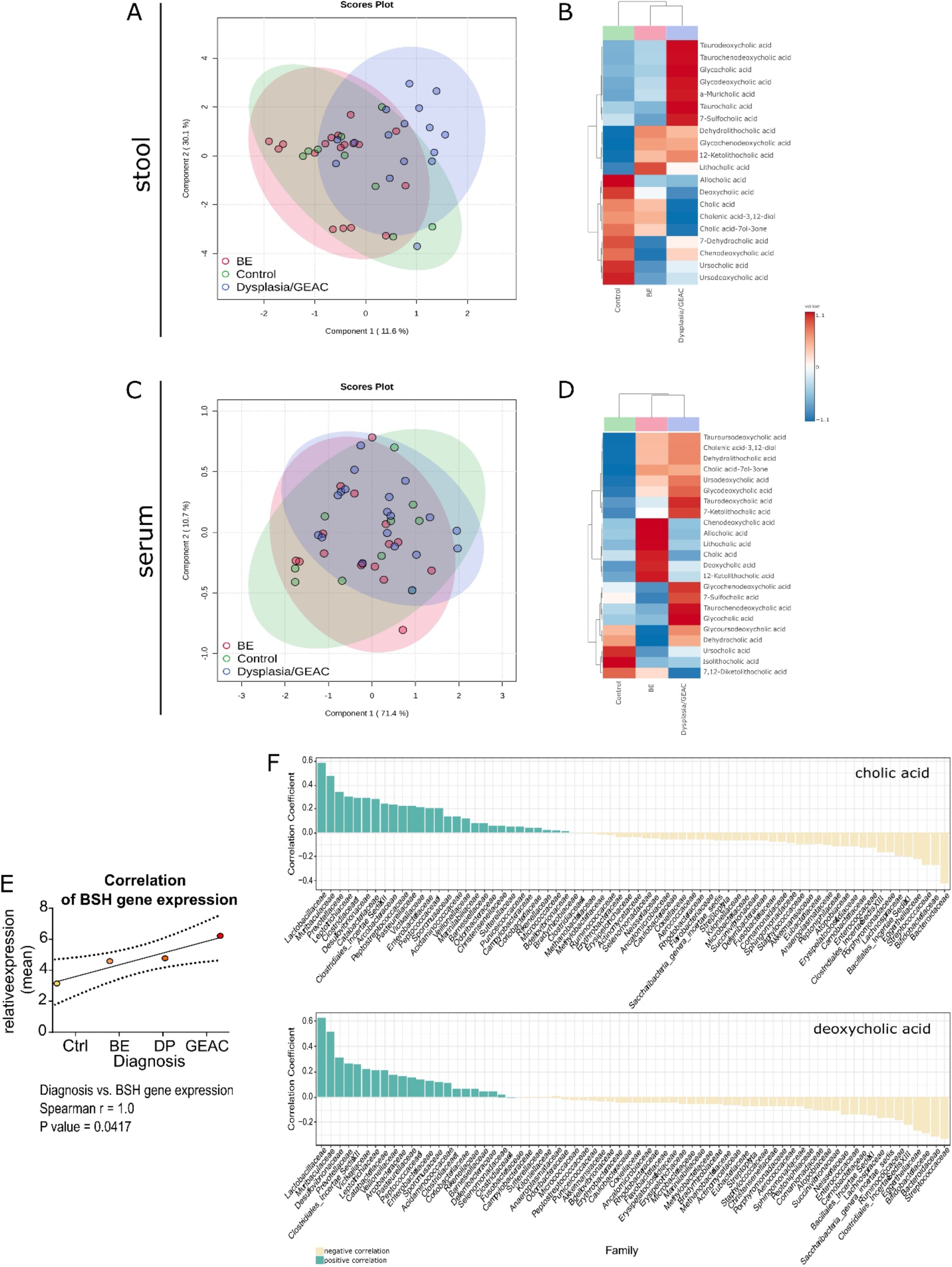
BA profile in stool and serum is linked to the disease stage in humans. **(A)** Partial least squares discriminant analysis (PLS-DA) of BA profiles in the stool of healthy controls (n=9), BE patients (n=18) and Dysplasia/GEAC patients (n=16). The BA profile of dysplastic and GEAC patients clustered differently than the healthy controls. **(B)** Clustering of the mean stool BA levels in healthy controls, BE and Dysplasia/GEAC patients shown as heatmap (Euclidean distance measure, ward clustering algorithm) **(C)** PLS-DA of BA profiles in the sera of healthy controls (n=10), BE patients (n=16) and Dysplasia/GEAC patients (n=19) showed no clear clusters **(D)** Clustering of the mean serum BA levels in human control-, BE- and GEAC patients shown as heatmap (Euclidean distance measure, ward clustering algorithm). **(E)** Correlation of gut bacterial BSH gene expression with the diagnostic stages of human patients. Spearman correlation analysis of the mean relative expression of BSH represented by delta Ct. **(F)** Bacteria families were ranked according to the correlation coefficients, which were calculated using the relative abundance of the bacteria family in all individuals and corresponding circulating CA and DCA levels. Only bacteria families with a correlation coefficient greater than 0.5 were considered. The family Lactobacillaceae correlated with levels of both, CA and DCA. (A-D) plots were created with data of targeted BA analysis. (B, D) Euclidean distance measure, ward clustering algorithm with Euclidean distance measure was applied for the heatmaps.

## DISCUSSION

Here we propose a novel concept that links GEAC carcinogenesis with alterations of the gut microbiota, enrichment of certain BA and associated expression and activation of the BA receptor FXR. We hypothesize that the increases in gut-borne BAs accelerate esophageal carcinogenesis via FXR-inhibition on progenitor cells at GEJ.

In L2-IL1B mice fed HFD, we observed an alteration of the gut microbiota with an associated shift in the BA profile of the gut, serum and BE tissue. The enrichment of cancer-promoting BAs (i.e. 12-Keto-LCA, TCDCA, Iso-DCA and TCA) (*27*) in the gut and most importantly, in the BE-tissue of HFD-fed mice, suggested these BAs to play a role during disease progression. In addition, numerous bacterial groups were negatively associated with inflammation and dysplasia while positively associated with metaplasia, implying that a diet-altered microbiota-metabolome-axis might represent a key factor that shapes the phenotypic fate of the progenitor cells at the GEJ towards differentiated goblet cells or dysplastic cells (*38, 39*).

In line with previous studies, we found upregulation of FXR in BE tissue of mice and patients, and downregulation associated with tumor development, suggesting that FXR activation might have a protective effect (*40, 41*). Indeed, genetic deficiency of FXR in L2-IL1B mice promoted and accelerated neoplasia. Loss of FXR led to formation of a tumor promoting microenvironment, with enrichment of pro-tumorigenic signaling pathways, less protective epithelial mucus production, and increased numbers of Lgr5 progenitor cells. Importantly, co-localization of Lgr5 and FXR in the BE region in L2-IL1B mice and enrichment of stem cell signaling pathways upon loss of FXR pointed to a direct function of FXR in Lgr5+ progenitor cells.

We hypothesize that a drop in FXR expression, together with non-physiological concentrations of BAs that inhibit FXR at the GEJ, could increase DNA damage and in turn, promote malignant transformation. Conversely, activation of FXR via its agonist OCA, altered the BA pool and showed anti-neoplastic effects. When examining FXR as a therapeutic target, we observed a protective effect of OCA in murine BE-organoids by attenuation of cytotoxic effects of the secondary BAs. When translating this into an *in vivo* proof-of principle, we observed that OCA reduced dysplasia and attenuated cell proliferation at the GEJ of HFD-fed L2-IL1B mice. Furthermore, the immune profile of OCA-treated HFD-fed mice was significantly altered, with reduced acute inflammation represented by neutrophil infiltration and elevated numbers of activated NKT cells. NKT cells have been shown to be controlled by a microbiota-associated BA-dependent cytokine signaling axis in a model of liver cancer (*42*), supporting the systemic immune-modulating BA effects in the current mouse model. In addition, OCA treatment resulted in decreased BSH expression in gut bacteria and reduced abundance of secondary BA producers such as Ruminococcaceae (*43*), pointing to diminished microbial potential for BA-transformation in OCA-treated mice.

Although fat metabolism and BA signatures differ between men a mice, complementary to our results in mice, BA levels and some associated microbial alterations in BE patients correlated with disease progression. However, the limited number of participants of this human cohort encourages more in depth-studies to confirm this clinical concept.

In summary, in a mouse model of BE and GEAC, a diet high in fat modified the composition of the intestinal microbiota, altered BA metabolism and levels in the gut, serum and distant tissue. Loss of FXR occurred with onset of tumorigenesis and resulted in a dysfunctional cellular capacity to compensate for BA-induced cell stress in progenitor cells at the GEJ. This led to establishment of a tumor-promoting microenvironment, characterized by dedifferentiation, progenitor cell expansion, DNA damage and increased inflammatory responsiveness, facilitating the dysplastic transformation of the epithelial progenitor cells. In mice, treatment with OCA reduced BA levels and ameliorated the dysplastic phenotype. In patients, BA profile in stool and serum correlated with disease progression. In conclusion, these data provide evidence for a novel mechanism by which a tightly interrelated diet-microbiome-metabolome axis accelerates tumorigenesis in the distal esophagus. Importantly, our results suggest that this acceleration of carcinogenesis can be prevented by treatment with the FXR agonist OCA. Similar therapies aimed to alter the systemic BA pool, including modifications to diet or directly to gut microbiome composition and function, may represent a novel avenue for preventing GEAC.

## Glossary

BA: Bile Acids
GEJ: Gastroesophageal Junction
BE: Barrett’s Esophagus
GEAC: Gastroesophageal Adenocarcinoma
HFD: High-Fat-Diet
DCA: Deoxycholic Acid
FXR: Farnesoid X Receptor
RXR: Retinoid X Receptor
OCA: Obeticholic Acid
L2-IL1B: mouse model of BE and GEAC
L2-IL1B-FXR KO: mouse model of BE and GEAC with whole-body knockout of FXR
CD: Control-Diet
LGD: Low-grade Dysplasia
HGD: High-grade Dysplasia
LCA: Lithocholic Acid
CDCA: Chenodeoxycholic Acid
TCA: Taurocholic Acid
GC: Goblet cell
NK: Natural Killer Cells
NKT: Natural Killer T Cells
BSH: Bile Salt Hydrolase
TLCA: Taurolithocholic acid
TDCA: Taurodeoxycholic acid
TCDCA: Taurochenodeoxycholic acid
GCA: Glycocholic acid
CA: Cholic Acid
TβMCA: Tauro-β-Muricholic Acid

## Funding

MQ, JA, HHW and TCW are supported by R01 CA272898

JA and MQ are supported by R01CA255298

MQ is supported bei Sander Stiftung and Deutsche Forschungsgesellschaft

H.H.W. is supported by funding from NSF (MCB-2025515), NIH (R01AI132403, R01DK118044, R01EB031935), Burroughs Wellcome Fund (PATH1016691), and the Irma T. Hirschl Trust.

H.H.W. is a scientific advisor of microbiome related companies SNIPR Biome, Kingdom Supercultures, Fitbiomics, Arranta Bio, Genus PLC, who were not involved in the study.

## List of Supplementary Materials

Materials and Methods: Breeding and mouse husbandry; Euthanasia, preparation, sample collection and disease evaluation; Inclusion criteria and sample collection for the BarrettNET study; Flow-cytometry for analysis of immune cells; Immunohistochemistry (IHC); In-Situ-Hybridization (ISH); RNA extraction and reverse transcription (RT); Quantitative Real Time PCR (qRT-PCR); Microarray analysis; High-throughput 16S rRNA gene amplicon-sequencing and analysis; DNA extraction and analysis of 16S rRNA sequencing data; OUT clustering and correlation analysis of the human stool samples; Mass spectrometry (MS) for targeted metabolomic analyses (sample preparation; performance, analysis and statistical evaluation of untargeted and targeted metabolomic analyses); Organoid culture from L2-IL1B mouse tissues (preparation of conditioned medium for organoid maintenance; isolation and maintenance of organoids); Organoid culture of human organoids; Treatment of organoids with bile acids; Click iT EdU flow cytometry assay kit for evaluation of cell proliferation of organoids; Histological analysis of organoids; Immunofluorescence staining of Paraffin-Embedded Organoid Slides; RNA extraction and downstream applications of organoids; DNA damage ELISA; Detailed statistical analysis.

Supplementary figures: Fig S1 to S7

Supplementary tables: Tables S1 to S2

## Notes

**Conflicts of interest:** The authors declare no potential conflicts of interest.

### Competing Interest Statement

The authors have declared no competing interest.

