## Supplementary Materials, Methods and Figures for "Microbiota metabolized Bile Acids accelerate Gastroesophageal Adenocarcinoma via FXR inhibition"

### SUPPLEMENTARY MATERIALS AND METHODS

#### Breeding and mouse husbandry

For microbiome and metabolome characterization of L2-IL1B mice on CD or HFD, L2-IL1B mice were backcrossed to C57BL/6J mice and bred and maintained under specific pathogen-free (SPF) conditions in the animal facility of the ZIEL Institute for Food and Health at the School of Life Sciences in Weihenstephan (WZW), TUM. L2-IL1B-FXR KO mice were bred and kept under SPF conditions in the animal facility of the Klinikum Rechts der Isar, TUM. For the OCA treatment studies in Germany, L2-IL1B mice were backcrossed to C57BL/6J mice and bred in a mouse facility from Charles River in Italy under SPF conditions. Following weaning and genotyping, mice were sent back to Germany at an age of 6-8 weeks. In Germany, all mice were kept under SPF conditions in the animal facility of the Klinikum Rechts der Isar, TUM. For the OCA treatment studies in New York, L2-IL1B mice were backcrossed to C57BL/6J mice and bred and maintained under SPF conditions in an animal facility of the Irving Cancer Research Center, Columbia University, NY.

#### Euthanasia, preparation, sample collection and disease evaluation, including mucus assessment at the GEJ

Mice were euthanized at specific timepoints and adequately dissected; feces, cecal content, blood from cardiac puncture and organs were subjected to downstream applications, including flow cytometry, FFPE, and storage at -80 °C for RNA-sequencing, 16S-sequencing and metabolomic analyses. Images taken from the stomach and esophagus during dissection were evaluated for tumor size and tumor coverage in percent using the software ImageJ(*1*). Macroscopic scores were determined as tumor size score and percentage of tumor coverage at the SCJ. (Criteria: tumor size score 0 = no abnormalities, 1= tumors < 0.5mm, 2= tumors <1mm, 3= tumors <2mm, 4= tumors <3mm). Mouse tissues were fixed in 4% paraformaldehyde and paraffin embedded, FFPE sections were cut and stained with H&E. Histopathologic scores were performed by an experienced Gastrointestinal Pathologist (DTP)-p following previously established criteria for the influx of immune cells per high-power field, metaplasia, and dysplasia (*2*). Inflammation was scored by the percentage of immune cells in a defined tissue area of the SCJ. Metaplasia was evaluated by the abundance of mucus production or cells per gland and the abundance of glands with mucus-producing cells in the BE area at the SCJ. Dysplasia was evaluated by the amount of cellular atypia and the presence of low- or high-grade dysplasia in single or multiple glands. Mucus production was assessed by periodic acid–Schiff (PAS) staining and quantified as percentage of PAS-positive area in BE regions.

#### Inclusion criteria and sample collection for the BarrettNET study

Inclusion criteria in the study included an age between 18-80 years, surveillance endoscopy in patients with already diagnosed BE without previously known occurrence of low-grade or high-grade dysplasia or GEAC and no presence of contraindications. Per patient and visit 4-6 endoscopic biopsies from the squamous epithelium of the cardia and BE regions, blood and fecal samples were collected. Biopsies (Preanalytics, PAXgene Tissue Container, Cat 765112) and blood (PAXgene blood DNA tubes 2,5 ml Cat 761165; Sarstedt S-Monovette® 5 ml 9NC, Cat. 05.1071) were collected in line with the endoscopy unit at the Klinikum Rechts der Isar, TUM, in Munich, Germany. Fecal samples were collected by the patients (Sarstedt, Stool Collection Tubes with Stool DNA Stabilizer, Cat. 1038111300). Biopsies were processed at the clinical pathology department according to the manufacturer’s protocol. Formalin-fixed and Paraffin-embedded (FFPE) samples were stored at -20 °C, stabilized blood samples were stored at -80°C or room temperature and stabilized fecal samples were stored at -80 °C under appropriate storage conditions.

#### Flow-cytometry for analysis of immune cells

Single-cell suspensions were generated as previously described(*3*). The following antibodies were used: APC-anti-F4/80, APC-e780-anti-cd11b1β, Alexa700-anti-Ly6G, eFluor450-lanti- CD45, PE-Ly6C, eFluor450-anti-CD4, APC-CD8a, FITC- anti-CD3, APCe780-NK1.1, and PE-anti-gamma delta TCR; 7-AAD was used to quantify live cells; all antibodies were purchased from eBioscience. Fluorescence-activating cell sorting (FACS) data were acquired on a Gallios flow cytometer (Beckman Coulter, Brea, CA) and analyzed using FlowJo software (FlowJo, Ashland, OR).

#### Immunohistochemistry (IHC)

IHC on sections from FFPE tissue was performed for detection of specific antigens in the tissue. Standard immunohistochemical procedures with citrate buffer antigen retrieval (H-3300, Vector Labs, Burlingame, CA) were conducted with the following antibodies: Ki67 (Abcam, AB15580; 1:1000, 4°C overnight), αSMA (Abcam, AB5694; 1:400, 4°C overnight), γH2AX (Cell Signaling, #9718; 1:750, 4°C overnight), rabbit-anti-mouse Caspase1 (1:50; 30min at room temperature) and TGR5 (Abcam, AB72608; 1:500, 4°C overnight). Quantification was assessed as the percentage of positive cells in the BE region, which were defined as the region between the squamous epithelium and the oxyntic mucosa of the stomach.

**In-Situ-Hybridization (ISH)**

ISH was done to detect antigen expression on the mRNA level for Lgr5 and FXR. The RNAscope 2.5 HD assay – Detection reagent BROWN (ACD) and all related reagents from ACD were used. The procedure was performed according to the manufacturer´s protocol using the Mm-Lgr5 target probe (ACD, Cat. No. 312171), the Hs-Lgr5 target probe (ACD, Cat. No 311021) and the Mm-NR1H4 target probe (ACD, Cat. No 484491) for detection of FXR expression. Quantification was assessed as the percentage of positive cells in the BE region as for IHC.

**RNA extraction and reverse transcription (RT)**

Tissue for RNA isolation was collected and stored overnight in 250 µl RNAlater™ (AM-7020; Invitrogen) at 4°C before long-term storage at -80°C. Isolation was performed using the RNeasy Mini Kit (74104; Qiagen) according to manufacturer's instructions. For tissue homogenization a SilentCrusher M (Heidolph) was employed. RNA was eluted in 20 µl PCR-grade water. RNA concentration and quality were measured on a Nano-Drop 2000 spectrophotometer (Thermo Scientific). RNA was directly subjected to reverse transcription (RT) or stored at -80°C until further use in downstream applications. For RNA extraction from stool, the Quick-RNA Fecal/Soil Microbe Microprep Kit (R2040, Zymo Research) was performed according to manufacturer's instructions. Reverse transcription was conducted with the QuantiTect Reverse Transcription Kit (205314, Qiagen) according to manufacturer´s instructions.

**Quantitative Real Time PCR (qRT-PCR)**

Target gene expression levels were evaluated by qRT-PCR on a LightCycler® 480 (Roche). PCR reactions were performed in a total volume of 10 µl per reaction using the QuantiFast SYBR Green PCR Kit (4000) (204057, Qiagen). For each reaction 10-25 ng RNA were applied in a volume of 1-2 µl, reactions were performed in triplicates. Glyceraldehyde 3-phosphate dehydrogenase (GAPDH), Cyclophilin A and beta-Actin served as standard housekeeping genes, 16S-RNA and bacterial GAPDH were used as housekeeping genes for determination of fecal bacterial gene expression. PCR conditions for all reactions were: 95°C for 3 minutes, followed by 40 cycles of 95°C for 30 seconds, 55°C for 30 seconds, and 72°C for 30 seconds. Primers and primer sequences were retrieved from the existing internal primer stock, from papers, collaborators, or designed via Primer-Blast (NCBI). All primer pairs were re-evaluated for self-complementation and gene specificity via blasting on Primer-Blast (NCBI). Primer amplification efficiencies were tested by generating a standard curve from PCR reactions of serial dilutions of the cDNA of positive control tissues. Furthermore, the melting curve of the primers was checked for quality control. Primer sequences of genes of interest and of housekeeping genes for fecal bacterial gene expression are listed in Table S1.

Table S1: Primers for quantitative Real Time - PCR

| Target | | Sequence | Melting T. | Notes |
|---|---|---|---|---|
| FXR | Forward | 5´-GGCTGCAAAGGTTTCTTCCG-3´ | 67.9 | |
| | Reverse | 5´-ACATTCAGCCAACATCCCCA-3´ | 67.6 | |
| IBABP | Forward | 5´-CACCATTGGCAAAGAATGTG-3´ | 65.7 | |
| | Reverse | 5´-AACTTGTCACCCACGACCTC-3´ | 66.7 | |
| SHP | Forward | 5´-AGCTGGGTCCCAAGGAGTAT-3´ | 63.7 | |
| | Reverse | 5´- CTTGAGGGTAGAGGCCATGA -3´ | 64.1 | |
| GAPDH | Forward | 5´-GACATCAAGAAGGTGGTGAAGCAG-3´ | 68 | |
| | Reverse | 5´-ATACCAGGAAATGAGCTTGACAAA-3´ | 64.5 | |
| βactin | Forward | 5´-CCGTGAACCCTAAGGCCAACC-3´ | 72.7 | |
| | Reverse | 5´-ACCCCGTCTCCGGAGTCCATC-3´ | 69.5 | |
| BSH | Forward | 5´-ATGGGCGGACTAGGATTACC-3´ | 63.8 | Working conc. 100 µM |
| | Reverse | 5´-TGCCACTCTCTGTCTTC-3´ | 54.2 | |
| 16S-RNA | Forward | 5´-TGATCCTGGCTCAGGACGAA-3´ | 68.3 | Working conc. 100 µM |
| | Reverse | 5´-TGCAAGCACCAATCAATACCA-3´ | 66.2 | |

**Microarray analysis**

Total RNA from SCJ and forestomach tissues from 12-month-old L2-IL1B WT and L2-IL1B-FXR KO mice (n=3) were extracted by TRIzol reagent (Invitrogen) according to the manufacturers protocol. Expression profiling was accomplished using Mouse gene 2.1 Affymetrix GeneChip® expression arrays. Differential expression was determined using Limma(*4*) as implemented in oneChannelGUI(*5*) operating as part of the Bioconductor

Suite(*6*) in the R statistical computing environment(*7*). Estimates of the statistical significance of overlap between gene sets were performed using the chi-square test[8] as implemented in R. Raw data have been deposited in the National Center for Biotechnology Information's Gene Expression Omnibus (GEO) (GSE103616). Functional annotation of phenotypes was performed by gene set enrichment analysis (GSEA) using the MSigDB database v5.2 where each phenotype was represented by three mice.

**High throughput 16S rRNA gene amplicon sequencing and analysis**

16S rRNA sequencing from patients' stool, saliva and PAXgene tissue samples was performed to characterize the gut, oral and local esophageal microbial microenvironment of the human patient cohort in different disease states. 16S rRNA gene sequencing from experimental mice on different diets and treatments was performed from cecal content and fecal samples to characterize the diet- and treatment-dependent microbial shifts in the mice. Therefore, DNA was extracted from the samples using different extraction methods.

**DNA extraction from fecal samples for 16S rRNA gene amplicon sequencing**

DNA from human fecal samples was extracted using a short modified version of the previously published Godon-protocol (*8*). Feces were already frozen with DNA stabilizer (Sarstedt, Stool Collection Tubes with Stool DNA Stabilizer, Cat. 1038111300). To 700 µl of the feces-DNA stabilizer mix, 250 µl 4M guanidinium thiocyanate and 500 µl 5% N-laurolylsarcosine were added. The probes were incubated for 1h at 70 °C. Sterile silica beads (0.1 mm, Biospec products) were used for bacterial cell lysis in a FastPrep-24 bead beater (MP biomedicals). Then, 15 mg polyvinylpyrrolidone was added, and the suspension was centrifuged at 15.000g at 4 °C for 3 min. The supernatant was collected and again centrifuged at 15.000g at 4 °C for 3 min. Thereafter, 500 µl of clear supernatant was collected, and 5 µl RNAse (10 mg/ml) were added to the samples followed by an incubation step for 20 min at 37 °C and shaking at 700 rpm. Subsequently, DNA was extracted with the NucleoSpin gDNA Clean-up kit following the manufacturer's protocol. DNA from human PAXgene biopsy samples was extracted using the PAXgene® tissue DNA-Kit (Preanalytix), following the manufacturer's protocol for purification of genomic DNA from sections of PAXgene-treated, paraffin-embedded tissue. For all extracted DNA samples, a sodium acetate precipitation was performed for purification and concentration of the samples. DNA samples were mixed 1:10 with a sodium acetate solution (3M). Then, 4 volumes of 100 % ethanol were added, mixed thoroughly, and

incubated over night at -20 °C. All samples were centrifuged at top speed for 30 min at 4 °C, and the supernatant was discarded. The pelleted DNA was washed with 500 µl ice-cold 80 % ethanol. The samples were centrifuged at top speed for 10 min at 4 °C, and the supernatant was discarded. If needed, the washing step was repeated once. The DNA pellet was air-dried and resuspended in 10 µL nuclease-free dH2O. After precipitation, concentration and purity of all samples was measured on a Nanodrop™ 1000 Spectrophotometer (Thermo Scientific). In a test-sequencing experiment, it was shown that DNA probes had sufficient quantitiy and quality for further analyses.
The V3/V4 region of 16S rRNA genes was amplified (25 cycles for fecal samples, 15 cycles for tissue biopsies) from 12 ng of metagenomic DNA using the bacteria-specific primers 341F and 785R following a two-step procedure to limit amplification bias(*9*). Amplicons were purified using the AMPure XP system (Beckmann), pooled in an equimolar amount, and sequenced in paired-end modus (PE275) using a MiSeq system (Illumina, Inc.) following the manufacturer's instructions.

**Analysis of 16S rRNA sequencing data**

16S rRNA sequencing data were analyzed using IMNGS, a web-based pipeline for processing of 16SRNA amplicon datasets(*10*), and RHEA, an R-based pipeline for data analysis and visualization. Beginning with the IMNGS workflow, resulting sequences were remultiplexed with a Perl script provided by the inventors of IMNGS termed remultiplexor(*10*). The IMNGS workflow itself is based on the UPARSE pipeline(*11*). In the IMNGS workflow, pairing, quality filtering and clustering of zero radius operational transcriptional units (zOTUs) were performed, wherefore USEARCH 8.0 was used(*12*). Therefore, all reads were trimmed to the position of the first base with a quality score smaller than three and then paired. The resulting sequences were size filtered and sequences with assembled size <300 and >600 nucleotides were excluded. Paired reads with an expected error bigger than three were also excluded. Remaining sequences were trimmed by five nucleotides on each side to avoid guanine-cytosine (GC) bias and nonrandom base composition. After processing of the remultiplexed data by the IMNGS workflow, a zOTU-table with associated sequences and taxonomic information for further analysis was generated. Additional analyses, including normalization, alpha- and beta-diversity, taxonomic abundance and correlation were performed using the Rhea-pipeline created for the R-interface RStudio(*13-15*).

**OTU clustering and correlation analysis of human stool samples**

Raw sequencing reads of the 16S-V4 region were analyzed using a previously described pipeline. Briefly, an OTU count per sample table was generated using USEARCH v11.0.667 (*12*) , and taxonomies were assigned using RDP classifier trained with 16S rRNA training set 18 (*16*). The samples with less than 3000 reads were filtered out. For each bacteria family, a Pearson correlation was calculated between its relative abundance and CA, DCA and TUDCAlevels across all samples.

**Mass spectrometry (MS) for targeted bile acid analysis**

Serum samples from the L2-IL1B and L2-IL1B-FXR KO mouse cohort, CD, HFD and HFD+OCA-treated L2-IL1B mice and healthy control individuals and patients diagnosed with BE, dysplasia and EAC from the BarrettNET study were used for metabolomic analyses. Also, cecal content and feces from CD, HFD and HFD+OCA-treated L2-IL1B mice and stool of patients were submitted for metabolomic analysis. Around 20 mg cecal and fecal/stool content or tissue were weighed in 2 ml bead beater tubes (CKMix 2 ml, Bertin Technologies, Montigny-le-Bretonneux, France) filled with ceramic beads (1.4 mm and 2.8 mm ceramic beads i.d.). Samples were later normalized to input weight.
For mouse samples, 1 ml methanol-based dehydrocholic acid extraction solvent (c=1.3 µmol/L) as an internal standard for work-up losses was added. For human samples, 100 mg of stool was extracted with 5 ml methanol-based dehydrocholic acid. Samples were homogenized using a bead beater (Precellys Evolution, Bertin Technologies) supplied with a Cryolys cooling module (Bertin Technologies, cooled with liquid nitrogen;3x20 seconds at 10.000rpm, 15 seconds breaks). The suspension was centrifuged (10 min, 8000 rpm, 10 °C) using an Eppendorf Centrifuge 5415R (Eppendorf, Hamburg, Germany). For metabolomic analysis of serum samples, 30 µL of serum was diluted with 270 µL methanolic dehydrocholic acid solution. After centrifugation as described above, the supernatant was used for analysis.
Targeted metabolomic analysis of bile acids was performed according to a method published by Reiter et al.(*22*). Briefly, 20 µl of isotopically labeled bile acids (ca. 7 µM each) were added to 100 µl of sample extract. Targeted bile acid measurement was done using a QTRAP 5500 triple quadrupole mass spectrometer (Sciex, Darmstadt, Germany) coupled to an ExionLC AD (Sciex, Darmstadt, Germany) ultrahigh performance liquid chromatography system. A multiple reaction monitoring (MRM) method was used for the detection and quantification of the bile acids. An electrospray ion voltage of −4500 V and the following ion source parameters were applied: curtain gas (35 psi), temperature (450 °C), gas 1 (55 psi), gas 2 (65 psi), and entrance potential (−10 V). The MS parameters and LC conditions were optimized using commercially available standards of

endogenous bile acids and deuterated bile acids, for the simultaneous quantification of selected 34 analytes. For separation of the analytes a 100 × 2.1 mm, 100 Å, 1.7 µm, Kinetex C18 column (Phenomenex, Aschaffenburg, Germany) was used. Chromatographic separation was performed with a constant flow rate of 0.4 ml/min using a mobile phase consisted of water (eluent A) and acetonitrile/water (95/5, v/v, eluent B), both containing 5 mM ammonium acetate and 0.1% formic acid. The gradient elution started with 25% B for 2 min, increased at 3.5 min to 27% B, in 2 min to 35% B, which was hold until 10 min, increased in 1 min to 43% B, held for 1 min, increased in 2 min to 58% B, held 3 min isocratically at 58% B. Then the concentration was increased to 65% at 17.5 min, with another increase to 80% B at 18 min, following an increase at 19 min to 100% B which was hold for 1 min., At 20.5 min, the column was equilibrated for 4.5 min at starting. The injection volume for all samples was 1 µL, the column oven temperature was set to 40 °C, and the auto-sampler was kept at 15 °C. Data acquisition and instrumental control were performed with Analyst 1.7 software (Sciex, Darmstadt, Germany). Targeted metabolomic analyses were analyzed as BA intensities, or respectively, if correlated to input weight and dilution, in nmol/g feces. Targeted Serum BA analysis comparing CD with HFD mice were visualized as heatmaps showing primary and secondary BA intensities between samples, as well as a correlation heatmap of BA levels with dysplasia scores and inversely with goblet cell ratios, representing the differentiation status. Bile acids detected via targeted BA analysis are listed in Table 2.

Table S2: Bile Acids measured in targeted BA metabolomic analysis

| | |
|---|---|
| 5β-Cholic acid-3α-ol-7-one,<br>7-Ketolithocholic acid, (7-KLCA) | 3-Dehydrocholic acid (3-DHCA) |
| 5β-Cholic acid-3α-ol-12-one,<br>12- Ketolithocholic acid, (12-KLCA) | 5β-Cholic acid-3α-ol-6,7-dione, (6,7-DKLCA) |
| α-Muricholic acid, (α-MCA) | 5β-Cholic acid-3α-ol-6-one, (6-KLCA) |
| β-Muricholic acid, (β-MCA) | 7-Dehydrocholic acid, (7-DHCA) |
| 5β-Cholen-24-oic acid-3,12-diol, Apocholic acid, (ApCA) | Cholic acid-7-sulphate, (7-SCA) |
| Chenodeoxycholic acid, (CDCA) | 12-Dehydrocholic acid, (12-DHCA) |
| Cholic acid, (CA) | 5β-Cholic acid-7α-ol-3-one, (Ca-7ol3one) |
| Deoxycholic acid, (DCA) | Glycocholic acid, (GCA) |

| | |
|---|---|
| Glycochenodeoxycholic acid, (GCDCA) | Glycoursodeoxycholic acid, (GUDCA) |
| Glycodeoxycholic acid, (GDCA) | Isolithocholic acid, (ILCA) |
| Hyodeoxycholic acid, (HDCA) | Taurocholic acid, (TCA) |
| Lithocholic acid, (LCA) | Taurohyodeoxycholic acid, (THDCA) |
| Tauro-α-Muricholic acid, (T-α-MCA) | Taurolithocholic acid, (TLCA) |
| Taurochenodeoxycholic acid, (TCDCA) | Tauro-ω-Muricholic acid, (T-ω-MCA) |
| Taurodeoxycholic acid, (TDCA) | Allocholic acid, (ACA) |
| Tauroursodeoxycholic acid, (TUDCA) | Ursocholic acid, (UCA) |
| Ursodeoxycholic acid, (UDCA) | 5β-Cholic acid-3α-ol-7,12-dione, (7,12-DKLCA) |
| Glycolithocholic acid, (GLCA) | Dehydrolithocholic acid, (DHLCA) |
| Murideoxycholic acid, (MDCA) | Glycohyocholic acid, (GHCA) |
| Allolithocholic acid, (ALCA) | Lithocholenic acid, (LCenA) |
| Glycohyodeoxycholic acid, (GHCA) | Obeticholic acid, (OCA) |
| Isodeoxycholic acid, (IDCA) | γ-Muricholic acid / Hyocholic acid, (γMCA) |

#### Analysis and statistical evaluation of targeted metabolomic analyses

For single BA analyses, values of the row data were first multiplied by 1000 and then log2 transformed. BA with N/A values were not considered. The data of all other BA was transferred to GraphPad for statistical analysis (details are listed below in the statistical part). For further analysis and visualization, including PLS-DA (Partial Least Squares – Discriminant Analysis), and clustered heatmap, the web-based tool metaboanalyst was employed (*23*).

### Organoid culture, maintenance and experiments

#### Preparation of conditioned medium for organoid maintenance

L-WRN (LWnt3A, R-spondin 3 and Noggin) conditioned medium with additional growth factors was used as growth medium for organoid culture. Media were conditioned wit L-

WRN-producing cells (ATCC® CRL3276™) according to the manufacturer´s protocol. The complete growth medium consisted of advanced DMEM/F12, supplemented with 10 % FBS, 1 % Penicillin/Streptomycin, HEPES buffer (pH 7.8) and Glutamax (all Gibco™ via ThermoFisher). For selection, G-418 and Hygromycin B (all Gibco™ via ThermoFisher) were used. For organoid culture, L-WRN conditioned media was supplemented with additional growth factors, including 1xB27 and N2 (17504044, 17502048, ThermoFisher), 50 ng/ml human EGF (AF-100-15, Peprotech) and 1mM N-acetyl cysteine (A7250, Sigma Aldrich).

#### Isolation and maintenance of murine organoids

Mouse organoid culture was performed according to the procedure published by Pastula et al. 2016 with minor adjustments(*24*). Resected and cleaned cardia tissue was cut into small pieces using surgical scissors and transferred to an Eppendorf tube containing 200-300 µl Accutase® cell detachment solution (A6964, Sigma-Aldrich). The tissue was incubated in Accutase® on a shaker for 15 min for enzymatic tissue digestion. Tissue pieces were transferred to a 50 ml collection tube containing 20 ml of ice-cold dPBS (14190144, Gibco™ via ThermoFisher) supplemented with 2 mM EDTA (AM9260G, Invitrogen™ via ThermoFisher) and EGTA (3054.2, Roth) each. All following steps were performed on ice. The tissue was incubated on a shaker on ice for 45 min for chemical tissue digestion. Supernatant was removed from sedimented tissue pieces, and tissue was washed and mechanically disintegrated by pipetting up and down in 10 ml of cold dPBS (14190144, Gibco™ via ThermoFisher) + 10 % FBS (10500064, Gibco™ via ThermoFisher). The tissue suspension was passed through a 70 µm cell strainer into a 50 ml tube. The tissue was removed from the strainer by washing with 10 ml of fresh dPBS + 10 % FBS. Mechanical disintegration, filtration, and recovery of the tissue from the strainer as described was repeated four times. The final volume of cell suspension was centrifuged at 4°C for 10 min at 400g. The supernatant was removed, and the cells were resuspended in 150-300 µl Matrigel® (354230, Corning®). Then, 50 µl ice-cold Matrigel® per well were plated in a 24-well plate prewarmed to 37°C and overlaid with 500 µl of complete growth media per well after solidification. The plate was incubated at 37 °C. Media was changed every 2-3 days, and organoids were passaged every 7-10 days depending on the growth rate. Media was removed, and Matrigel® disrupted by repetitive pipetting with ice-cold dPBS + 10 % FBS buffer, and the suspension from each well was pooled into a 15 ml collection tube. The washing step was repeated. Matrigel® and organoids were gently disrupted by repetitive pipetting of the suspension in the 15 ml collection tube. The tube was then centrifuged at 4°C for 10 min. at 400g. Afterwards, the supernatant and Matrigel® debris were carefully removed. The cell pellet was resuspended in fresh Matrigel®, plated into a prewarmed 24 well plate and overlaid with media as previously described. Wells were expanded 1:2-1:4 per passage depending on the growth rate of the organoids.

#### Isolation and maintenance of human organoids

Organoids were isolated from biopsies taken from BE- and GEAC-patients. Prior to biopsy collection, explicit informed consent was obtained from all patients in accordance with institutional guidelines and ethical standards (FREEZE-BiObank).

Freshly taken biopsies in tissue storage solution (Miltenyi Biotec) + 10 µM Y-27632/Rock Inhibitor were transferred and digested in a FALCON containing prewarmed 10mL digestion medium (DMEM+++ composed of advanced DMEM/F12, 1 % Penicillin/Streptomycin, 1% HEPES buffer and 1% Glutamax, together with 1% Primocin, 1 mg of Collagenase XI 1 and 10mg of Dispase II). The biopsy was incubated for 30-90 minutes in a water bath (37 °C) and vigorously shaken to promote tissue disruption and release of glands and crypts during digestion. After incubation step, FALCON was centrifuged (5min, 400g, RT), supernatant was discarded, 15 mL digestion buffer were added and sample was centrifuged again. This washing step was repeated for three times. After the last washing step, tissue pellets were resuspended in 100% MG and plated as 20µL domes. After the solidification of domes for 10 minutes, the medium (DMEM+++ containing 1xB27, 1xN2, 50µg/mL human EGF, 1mM N-Acetylcystein, 10mM Nicotinamide, 5µM P38 MAPK Inhibitor, 2µM TGF beta inhibitor, 0.5µM CHIR99021, 10nM Gastrin, 100ng/mL FGF-10, 10µM Rock-Inhibitor, 100µg/mL Primocin) was added.

#### Treatment of organoids with bile acids

Mouse cardia organoids were isolated from L2-IL1B mice. Treatment was started two days after the $3^{rd}$ - $4^{th}$ passage of the organoids. The organoid treatment was performed for 72 h with OCA, DCA or TβMCA or OCA + DCA or OCA + TβMCA in concentrations of 10 and 100 µM diluted in the medium, respectively. Every 24 hrs, cell numbers were counted and cultures photographed, and medium was changed subsequently. After 72h, 2-3 wells with organoids were pooled, and RNA was extracted using the RNeasy Micro Kit (Qiagen, Germany), and conditioned media from every treatment condition was collected and stored. The number of organoids counted every 24h was evaluated as percentage of organoid numbers relative to 0h.

#### Click iT EdU flow cytometry assay kit for evaluation of cell proliferation of organoids

Cell proliferation rates in treated organoids were evaluated using the Click-iT™ EdU Alexa Fluor™ 488 Flow Cytometry Assay Kit (C10425, Invitrogen™ via ThermoFisher). Reagent were prepared and cells labeled, fixed and permeabilized according to the manufacturer´s protocol with minor modifications. Organoid cells from approximately 6 wells per treatment group were treated with 50 µM EdU in fresh conditioned medium for two hours in an incubator at 37 °C. After two hours, organoids were harvested with 500 µl of ice-cold dPBS + 10 % FBS per well, pooled and centrifuged for 10 min at 4 °C and 400 rcf. The cell pellet was then incubated in 1 ml of 0.25 % Trypsin-EDTA (25200056, Gibco™ via ThermoFisher) for 15 min on a rotation wheel in an incubator at 37 °C with occasional vortexing to obtain single cells. The reaction was stopped with dPBS + 10 % FBS, the cells were again centrifuged, and the supernatant was discarded. After single cell isolation, cells were washed with 3 ml of 1% BSA (9418, Sigma) in dPBS and centrifuged as described before. The supernatant was discarded, and the pellet was thoroughly resuspended in 100 µl of Click-iT® fixative. Cells were incubated for 15 min at room temperature (RT) on a rotation wheel in the dark. Cells were washed with 3 ml of 1% BSA in dPBS and pelleted by centrifugation as described before. The cell pellet was resuspended in 100 µl of 1X Click-iT® saponin-based permeabilization and wash reagent. The Click-iT® EdU reaction cocktail was prepared according to the manufacturer´s protocol. Then, 0.5 ml of freshly prepared reaction cocktail were added to the organoid cells in 1X Click-iT® saponin-based permeabilization and wash reagent, and samples were incubated for 30 min. at RT on a rotation wheel in the dark Cells were again washed with 3 ml of 1% BSA in dPBS and pelleted by centrifugation as above. The supernatant was removed, and the cells were resuspended in 100 µl of 1X Click-iT® saponin-based permeabilization and wash reagent. Cells were stained for DNA content and analyzed on a Gallios flow cytometer (Beckman Coulter), using an excitation laser at 488nm, and measuring emission using a green bandpass emission filter (530/30 nm). Results were analyzed using FlowJo software (TreeStar).

#### Histological analysis of organoids

For fixation and embedding of organoids for FFPE sections, organoids were grown on cover slips placed into a 24-well plate. Organoids were washed three times with 500 µl of PBS$^{+}$ (dPBS + 0.9mM $CaCl_2$ and 0.493 mM of $MgCl_2$) per well for one minute before fixation in 500 µl of 4 % PFA for 30 min at RT on a shaker. After fixation, washing steps were repeated. Organoids were transferred to Bio-Net histology cassettes (09-0403, Langenbrinck GmbH), dehydrated, paraffin-embedded and cut into sections using a manual microtome. Staining and IHC were performed as for tissue sections. Quantification was assessed as the number of positive cells compared to all cells per organoid. All organoids from one mouse, which had been subjected with the same treatment were counted as technical replicates. Organoids from different mice were defined as biological replicates. Organoids from n=3 mice per condition were used for statistical analysis. To this end, ordinary one-way ANOVA with Tukey test to correct for multiple comparisons was performed.

**Immunofluorescence Staining of Paraffin-Embedded Organoid Slides**

Human organoids samples (3 GEAC, 3BE) were fixed after 2-4 passages using 4% paraformaldehyde solution (PFA) in phosphate-buffered saline (PBS) with a pH of 7.4 (sc-281692, ChemCruz via Santa Cruz Biotechnology) for 30 minutes at the room temperature. Fixed organoids were dehydrated using graded ethanol series and xylene, and were finally embedded in paraffin blocks using a standard tissue processing protocol.

FFPE organoid blocks were cut into 3.0-µm-thick sections using a microtome. Sections were mounted onto positively charged glass slides and dried at room temperature overnight. Slides were immersed in xylene for 15 minutes (2 times) to remove paraffin. Sections were rehydrated through a series of graded ethanol solutions (100%, 95%, and 70%). Slides were subjected to heat-mediated antigen retrieval using a target retrieval solution, pH 9.0 (S2367, Dako) in a pressure cooker for 15 minutes. After cooling, slides were washed with 1x wash buffer (S3006, Dako).Then slides were blocked with 10% goat serum (ENG9010-10, BIOZOL Diagnostica Vertrieb GmbH), 1: 100 Fc block anti CD16/CD32 (553142, BD Biosciences) in antibody diluent with background reducing components (S3022, Dako) for 1 hour at room temperature to minimize nonspecific binding. Next, sections were incubated with the primary antibody to FXR (sc-13063, Santa Cruz Biotechnology) in antibody diluent (1:25) with background reducing compounds overnight at 4°C in a humidified chamber. Slides were washed three times for 5 minutes each with 1x wash buffer to remove unbound primary antibodies.

Fluorescent-labeled secondary antibodies 1:400 (A11034, Invitrogen) were applied to the sections for 1 hour at room temperature, then slides were washed 3 times more. To minimize autofluorescence of samples, we used Quenching Kit (SP-8500, Vector Laboratories Inc.). As a final step of this kit, slides were mounted using an anti-fade medium with DAPI to preserve fluorescence signals and counterstain nuclei.

Per one sample 5 different fields of view (20x magnification) containing organoids were taken. Images were analyzed with CellProfiler 4.2.5*. The nuclear signal of FXR was measured in the regions of interests defined by the DAPI signal. Nuclei were classified as positive if their mean FXR intensity exceeded that of 75% of all nuclei. The result was expressed as the percentage of positive nuclei relative to the total number of nuclei in field of view.

#### RNA extraction and downstream applications

In brief, 3-6 wells of organoids were harvested with 500 µl of dPBS + 10 % FBS per well, cells were pooled and centrifuged for 10 min at 4 °C and 400 rcf. The supernatant was removed, and the pellet was resuspended in 500 µl collagenase/ dispase (1 mg/ml) and incubated at 37 °C for 1 hour to digest Matrigel® residuals enzymatically. The reaction was stopped with 500 µl 5 mM EDTA (AM9260G, Invitrogen™ via ThermoFisher) in dPBS. The tube was filled 10 ml with dPBS, centrifuged for 10 min at 4 °C and 400 rcf, and the supernatant was removed. The remaining cell pellet was lysed in 600 µl RLT-buffer (Lysis buffer; 1015762; Qiagen) supplemented with 1 % beta-mercaptoethanol (4227.3, Roth). RNA isolation was performed using the RNeasy Mini Kit (74104; Qiagen) according to the manufacturer´s protocol. Reverse transcription PCR and qRT-PCR were both conducted as described for tissue samples.

#### DNA damage ELISA

OxiSelect Oxidative DNA Damage ELISA, 8´OhdG Quantitation (Cell Biolabs, Inc. STA-320) was performed according to the manufacturer's protocol using conditioned media from the different treatments done in the organoid cultures. Conditioned media from 12m-old L2-IL1b mouse organoids treated with OCA, DCA, TβMCA, OCA+DCA or OCA+TβMCA and without treatment were analyzed after 48h and 72h treatment, respectively. Data was acquired on a multiskan FC microplate reader (Thermo Scientific) and analyzed using Microsoft Excel and GraphPad Prism version 8.

**SUPPLEMENTARY FIGURES:**

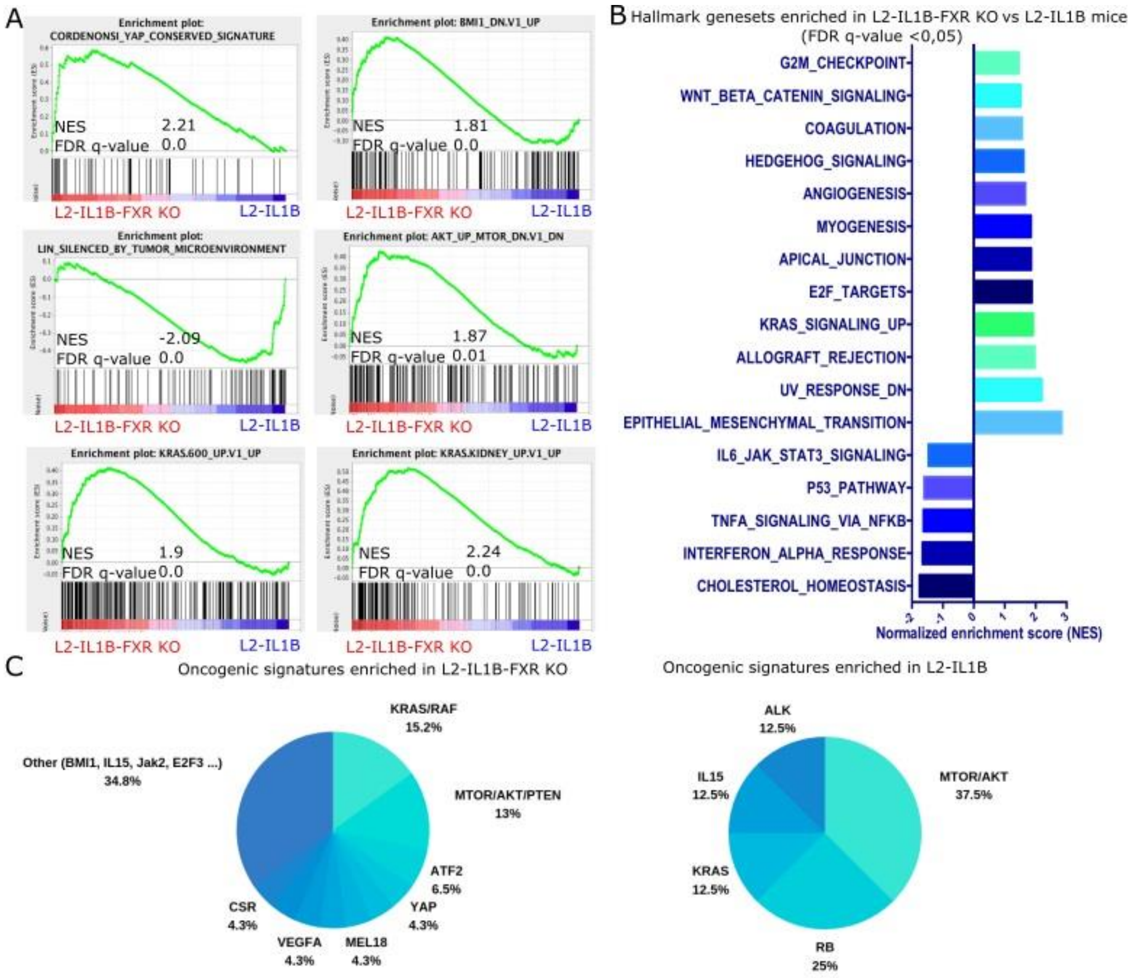


**Supplementary Figure 1: Microarray analysis shows enrichment of pro-tumorigenic pathways and genes in L2-IL1B FXR-KO compared to L2-IL1B littermates**

- **(A)** Gene set enrichment analysis (GSEA) showed enrichment of YAP, BMI1, AKT and KRAS signaling pathways, as well as the increased epigenetically induced silencing of the CST6 tumor suppressor gene in L2-IL1B-FXR KO compared to L2-IL1B mice (gene set enrichment analysis of cardia tissue microarray; n=3).
- **(B)** Significantly enriched hallmark gene sets in L2-IL1B-FXR KO compared to L2-IL1B mice (FDR q-value ≤ 0.05).
- **(C)** Oncogenic signature pathways enriched in L2-IL1B-FXR KO (n=18) compared to L2-IL1B (n=16) mice.

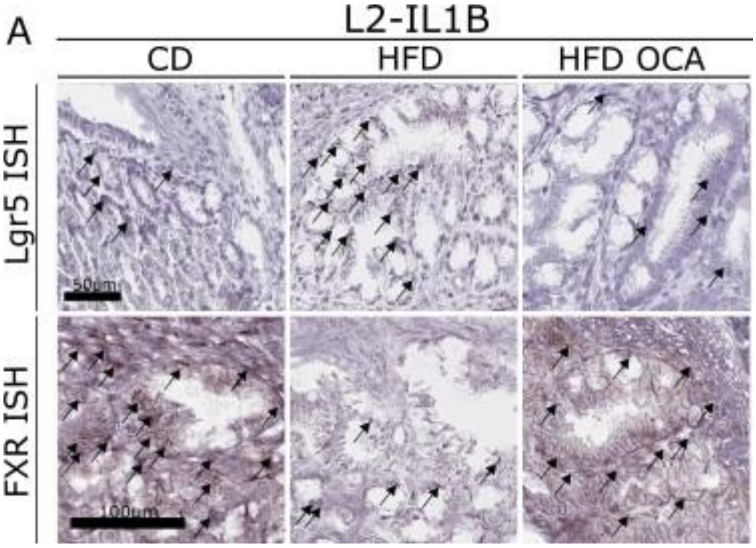


**Supplementary Figure 2: Scoring and analysis of FXR expression in L2-IL1B mice treated with CD, HFD and HFD+OCA**

- **(A)** Inflammation and metaplasia scores of 6-9-month-old L2-IL1B mice treated with CD, HFD or HFD+OCA, respectively, showed no significant differences between treatments (n=7-12; $p \geq 0.05$).
- **(B)** Representative pictures of Lgr5 ISH of 9-month-old L2-IL1B mice treated with CD, HFD or HFD+OCA.

(A) Data are depicted as mean with SD. For statistical evaluation of A, Kruskal-Wallis tests with Dunn´s test for correction of multiple comparisons were performed.

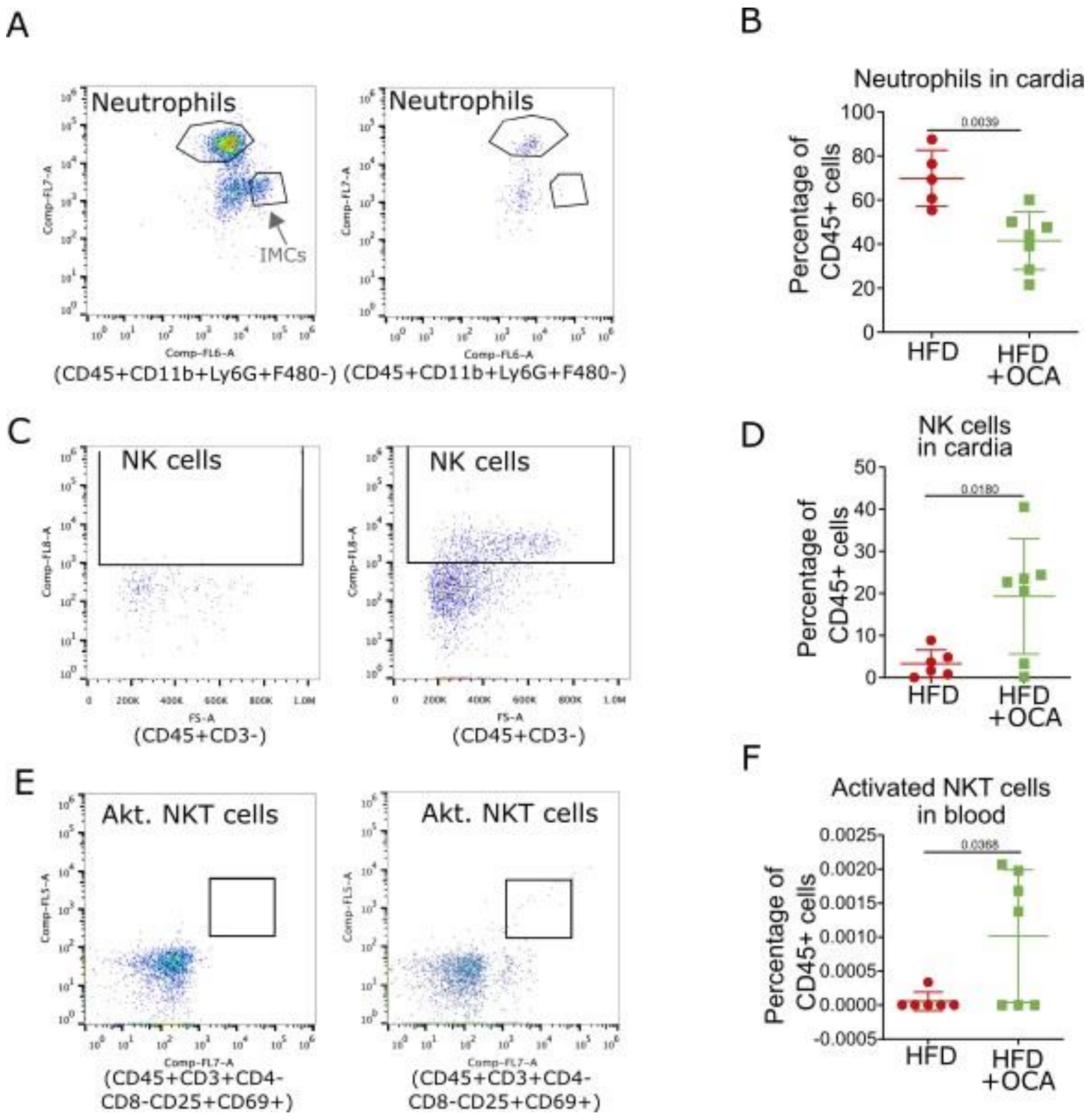


**Supplementary Figure 3: HFD OCA mice display altered numbers of neutrophils, NK and activated NKT cells**

- **(A)** Representative gating schemes of neutrophils in the cardia of L2-IL1B HFD-fed and HFD+OCA-fed mice
- **(B)** The percentage of neutrophils in the cardia region of HFD+OCA-fed compared to HFD-fed mice significantly decreased (n=6-7).
- **(C)** Representative gating schemes of NK cells in the cardia of L2-IL1B HFD-fed and HFD+OCA-fed mice
- **(D)** The percentage of NK cells in the cardia region of HFD+OCA-fed compared to HFD-fed mice significantly increased (n=6-7).
- **(E)** Representative gating schemes of NKT cells in the blood of L2-IL1B HFD-fed and HFD+OCA-fed mice

**(F)** The percentage of activated NKT cells in the blood of HFD+OCA-fed compared to HFD-fed mice significantly increased (n=6-7).

For (B, D, F) data are presented as mean percentage of CD45+ cells with SD. For statistical evaluation, unpaired, two-tailed t-tests were performed.

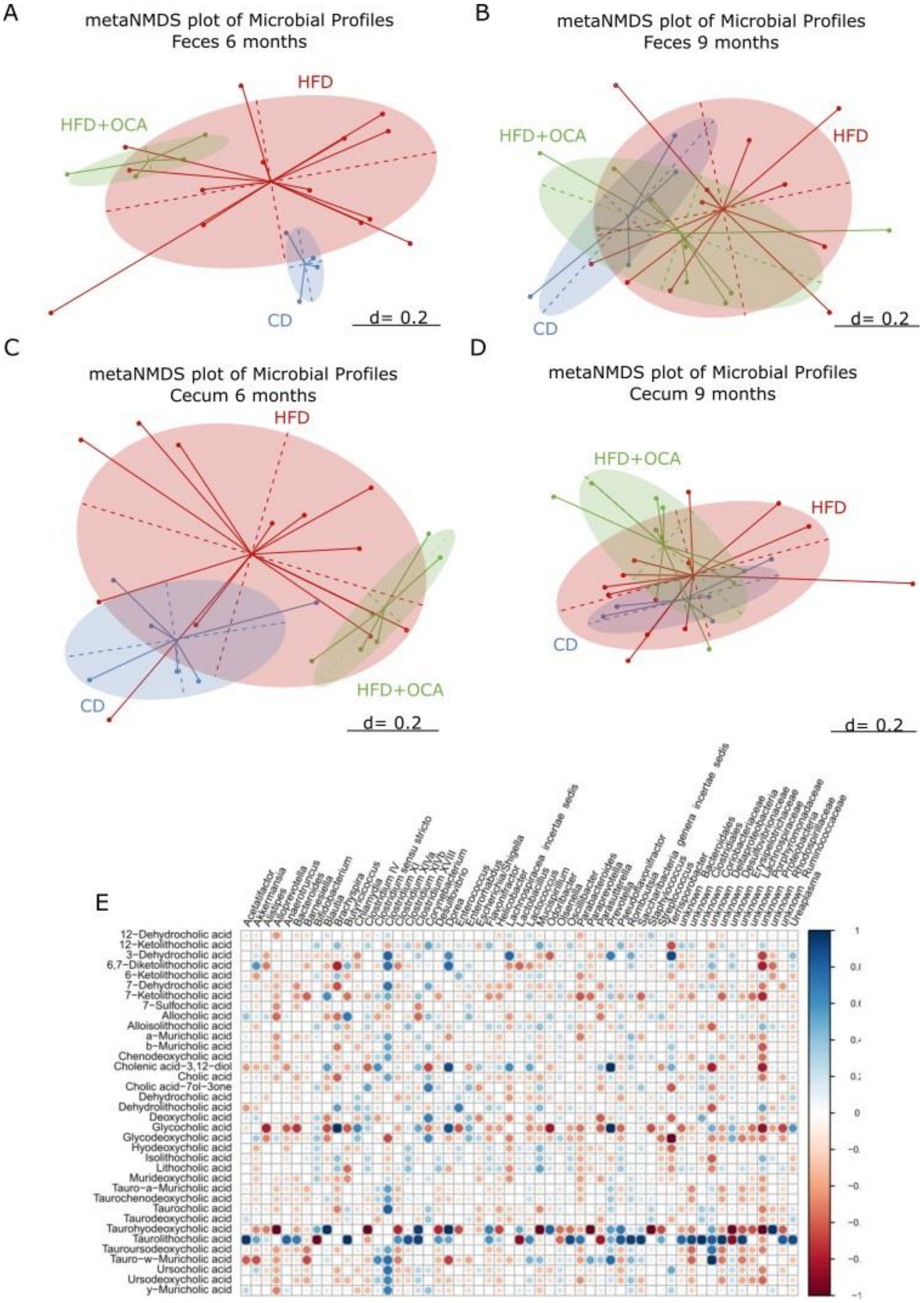


**Supplementary Figure 4: Beta diversity of fecal and cecal microbial profiles and correlation analysis between fecal BA levels and microbial genera in L2-IL1B mice.**

**(A)** Meta–Normalized Multidimensional Scaling (metaNMDS) plot of feces from 6-month-old L2-IL1B mice fed with CD, HFD or HFD+OCA showed significant separation of study groups in terms of bacterial community composition (p=0.002)

**(B)** metaNMDS plot of feces from 9-month-old L2-IL1B mice fed with CD, HFD or HFD+OCA showed no significant separation of study groups in terms of bacterial community composition (p=0.214).

**(C)** metaNMDS plot of cecum from 6-month-old L2-IL1B mice fed with CD, HFD or HFD+OCA showed significant separation in terms of bacterial community composition (p=0.002).

**(D)** metaNMDS plot of cecum from 9-month-old L2-IL1B mice fed with CD, HFD or HFD+OCA showed no significant separation of study groups in terms of bacterial community composition (p=0.289).

**(E)** Correlation analysis between fecal BA levels and the abundance of fecal microbial genera in L2-IL1B mice fed with CD, HFD or HFD+OCA.

For (A-D) Permutational Multivariate Analysis of Variance (PERMANOVA) test was used for statistical comparison. The microbiota phylogenetic distances were evaluated through the generalized UniFrac distance. Each point represents the microbiota composition of one sample. (E) Correlation analyses were performed using the Rhea pipeline for 16S rRNA-sequencing data.

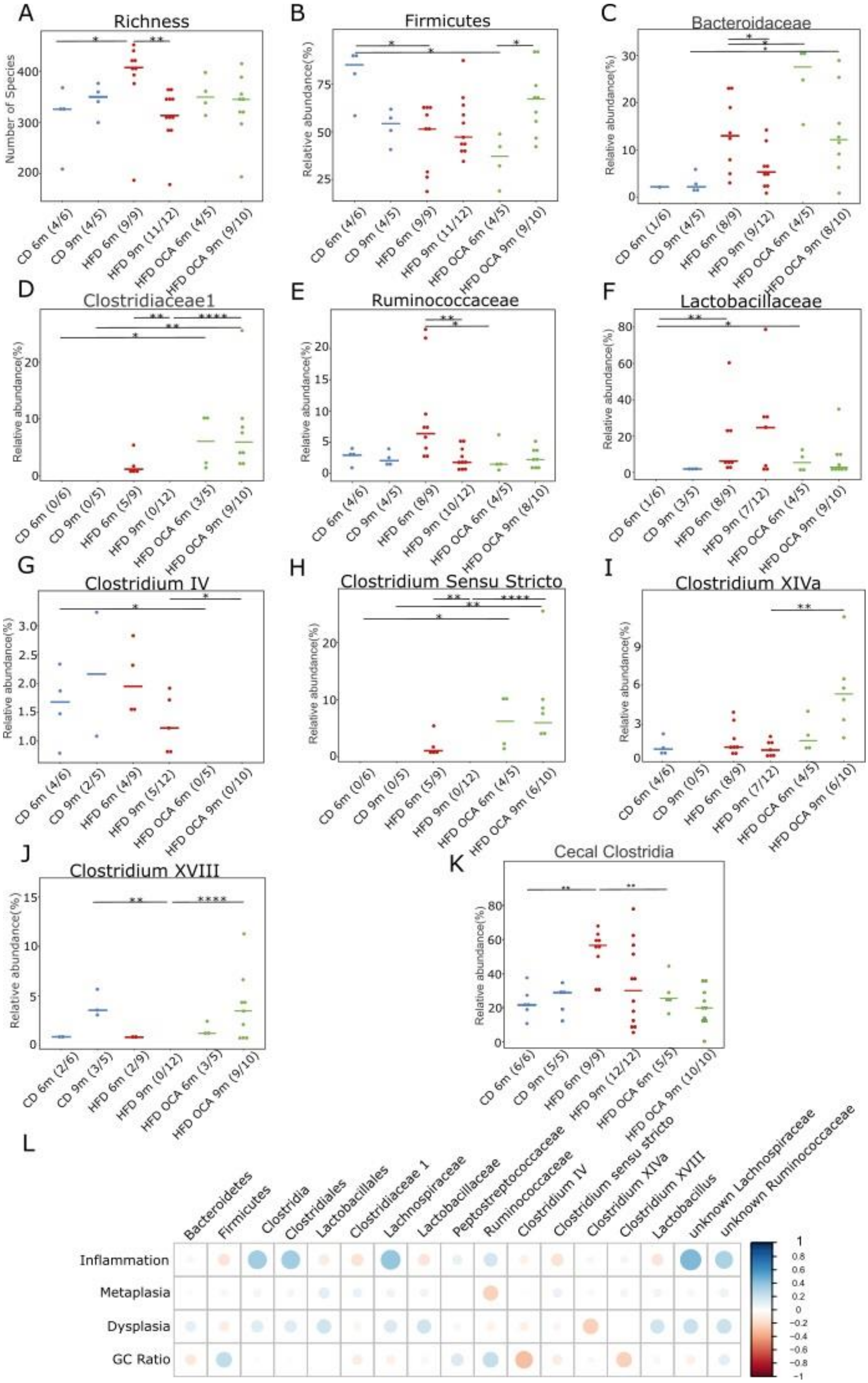

A
Richness
Number of Species
CD 6m (4/6)
CD 9m (4/5)
HFD 6m (9/9)
HFD 9m (11/12)
HFD OCA 6m (4/5)
HFD OCA 9m (9/10)
B
Firmicutes
Relative abundance(%)
CD 6m (4/6)
CD 9m (4/5)
HFD 6m (9/9)
HFD 9m (11/12)
HFD OCA 6m (4/5)
HFD OCA 9m (9/10)
C
Bacteroidaceae
Relative abundance(%)
CD 6m (1/6)
CD 9m (4/5)
HFD 6m (8/9)
HFD 9m (9/12)
HFD OCA 6m (4/5)
HFD OCA 9m (8/10)
D
Clostridiaceae1
Relative abundance(%)
CD 6m (0/6)
CD 9m (0/5)
HFD 6m (5/9)
HFD 9m (0/12)
HFD OCA 6m (3/5)
HFD OCA 9m (9/10)
E
Ruminococcaceae
Relative abundance(%)
CD 6m (4/6)
CD 9m (4/5)
HFD 6m (8/9)
HFD 9m (10/12)
HFD OCA 6m (4/5)
HFD OCA 9m (8/10)
F
Lactobacillaceae
Relative abundance(%)
CD 6m (1/6)
CD 9m (3/5)
HFD 6m (8/9)
HFD 9m (7/12)
HFD OCA 6m (4/5)
HFD OCA 9m (9/10)
G
Clostridium IV
Relative abundance(%)
CD 6m (4/6)
CD 9m (2/5)
HFD 6m (4/9)
HFD 9m (5/12)
HFD OCA 6m (0/5)
HFD OCA 9m (0/10)
H
Clostridium Sensu Stricto
Relative abundance(%)
CD 6m (0/6)
CD 9m (0/5)
HFD 6m (5/9)
HFD 9m (0/12)
HFD OCA 6m (4/5)
HFD OCA 9m (6/10)
I
Clostridium XIVa
Relative abundance(%)
CD 6m (4/6)
CD 9m (0/5)
HFD 6m (8/9)
HFD 9m (7/12)
HFD OCA 6m (4/5)
HFD OCA 9m (6/10)
J
Clostridium XVIII
Relative abundance(%)
CD 6m (2/6)
CD 9m (3/5)
HFD 6m (2/9)
HFD 9m (0/12)
HFD OCA 6m (3/5)
HFD OCA 9m (9/10)
K
Cecal Clostridia
Relative abundance(%)
CD 6m (6/6)
CD 9m (5/5)
HFD 6m (9/9)
HFD 9m (12/12)
HFD OCA 6m (5/5)
HFD OCA 9m (10/10)
L
Bacteroidetes
Firmicutes
Clostridia
Clostridiales
Lactobacillales
Clostridiaceae 1
Lachnospiraceae
Lactobacillaceae
Peptostreptococcaceae
Ruminococcaceae
Clostridium IV
Clostridium sensu stricto
Clostridium XIVa
Clostridium XVIII
Lactobacillus
unknown Lachnospiraceae
unknown Ruminococcaceae
Inflammation
Metaplasia
Dysplasia
GC Ratio

**Supplementary Figure 5: Treatment with obeticholic acid (OCA) changes the BA-metabolizing gut microbiome in L2-IL1B mice**

Only relevant significant changes are shown.

- **(A)** Bacterial community richness increased in the feces of 6-m-old HFD-fed mice compared to CD-fed mice (p=0.0336). While the richness in both CD and HFD+OCA diets remained constant between 6 month and 9 month of age, richness significantly decreased within the HFD cohort (HFD 6m vs HFD 9m p=0.0016).
- **(B)** The relative abundance of the phylum Firmicutes decreased in both 6m old HFD-fed (p= 0.0336) and HFD+OCA-fed (p= 0.0286) mice compared to CD-fed mice and increased in 9m old compared to 6m old HFD+OCA-fed mice (p=0.0196).
- **(C)** The relative abundance of the family Bacteroidaceae significantly increased in 6-m-old HFD+OCA-fed compared to HFD-fed and in 9-m-old HFD+OCA-fed compared to CD-fed mice (CD 9m vs HFD+OCA 9m p=0.0485; HFD 6m vs HFD+OCA 6m p=0.0283), while the abundance decreased in HFD-fed mice from 6 to 9 months of age (HFD 6m vs HFD 9m p=0.0464).
- **(D)** The family *Clostridiaceae 1* were only abundant in 6-m-old HFD-fed and 6-m- and 9-m-old HFD+OCA-fed mice (CD 6m vs HFD+OCA 6m p=0.0286; CD 9m vs HFD+OCA 9m p=0.0070; HFD 9m vs HFD+OCA 9m p=0.0001; HFD 6m vs HFD 9m p=0.0081).
- **(E)** The relative abundance of the family of *Ruminococcaceae* decreased in both 9-m-old HFD-fed and 6-m-old HFD+OCA-fed mice compared to 6-m-old HFD-fed mice (HFD 6m vs HFD 9m p=0.0044; HFD 6m vs HFD+OCA 6m p=0.0485).
- **(F)** The family of *Lactobacillaceae* was significantly higher abundant in 6-m-old HFD-fed and HFD+OCA-fed compared to 6-m-old CD-fed mice (CD 6m vs HFD 6m p=0.0070; CD 6m vs HFD+OCA 6m p=0.0286).
- **(G)** The genus *Clostridium* cluster IV was only abundant in CD- and HFD-fed mice (CD 6m vs HFD+OCA 6m p=0.0286; HFD 9m vs HFD+OCA 9m p=0.0379).
- **(H)** The genus *Clostridium sensu stricto* was significantly higher abundant compared to 6-m- and 9-m-old CD-fed mice and compared to 9-m-old HFD-fed mice (CD 6m vs. HFD+OCA 6m p=0.0286; CD 9m vs. HFD+OCA 9m p=0.0070; HFD 9m vs.HFD+OCA 9m p=0.0001). Abundance droped in HFD-fed mice from 6 to 9 months of age (p=0.0081).

**(I)** The *Clostridium cluster XIV* was significantly more abundant in 9m old HFD+OCA-fed mice compared to HFD-fed mice (p=0.0023).

**(J)** The genus *Clostridium XVIII* was only abundant in 6-m-old HFD-fed and in 6-m- and 9-m-old HFD+OCA-fed mice (CD 9m vs HFD 9m p=0.0088; HFD 9m vs HFD+OCA 9m p<0.0001).

**(K)** In the cecal content of the mice, the class of Clostridia was significantly enriched in abundance in 6-m-old HFD-fed compared to CD- and HFD+OCA-fed mice (HFD 6m vs CD 6m p=0.0016; HFD 6m vs HFD+OCA 6m p=0.0040).

**(L)** Correlation analysis between the abundance of bacterial families with BA metabolizing capacities in all treatment groups vs inflammation, metaplasia, and dysplasia scores as well as goblet cell (GC) ratio. Correlation analyses were performed using the Rhea pipeline for 16s-sequencing data. Clostridia, Lactobacillales and *Lachnospiraceae* and unknown *Ruminococcaceae* were the main microbial groups positively correlated with dysplasia scores.

For (A-K) relative abundances of microbiota are represented as mean. For statistical analysis, pairwise Wilcoxon rank sum test or Fisher´s exact test, if appropriate, were used.

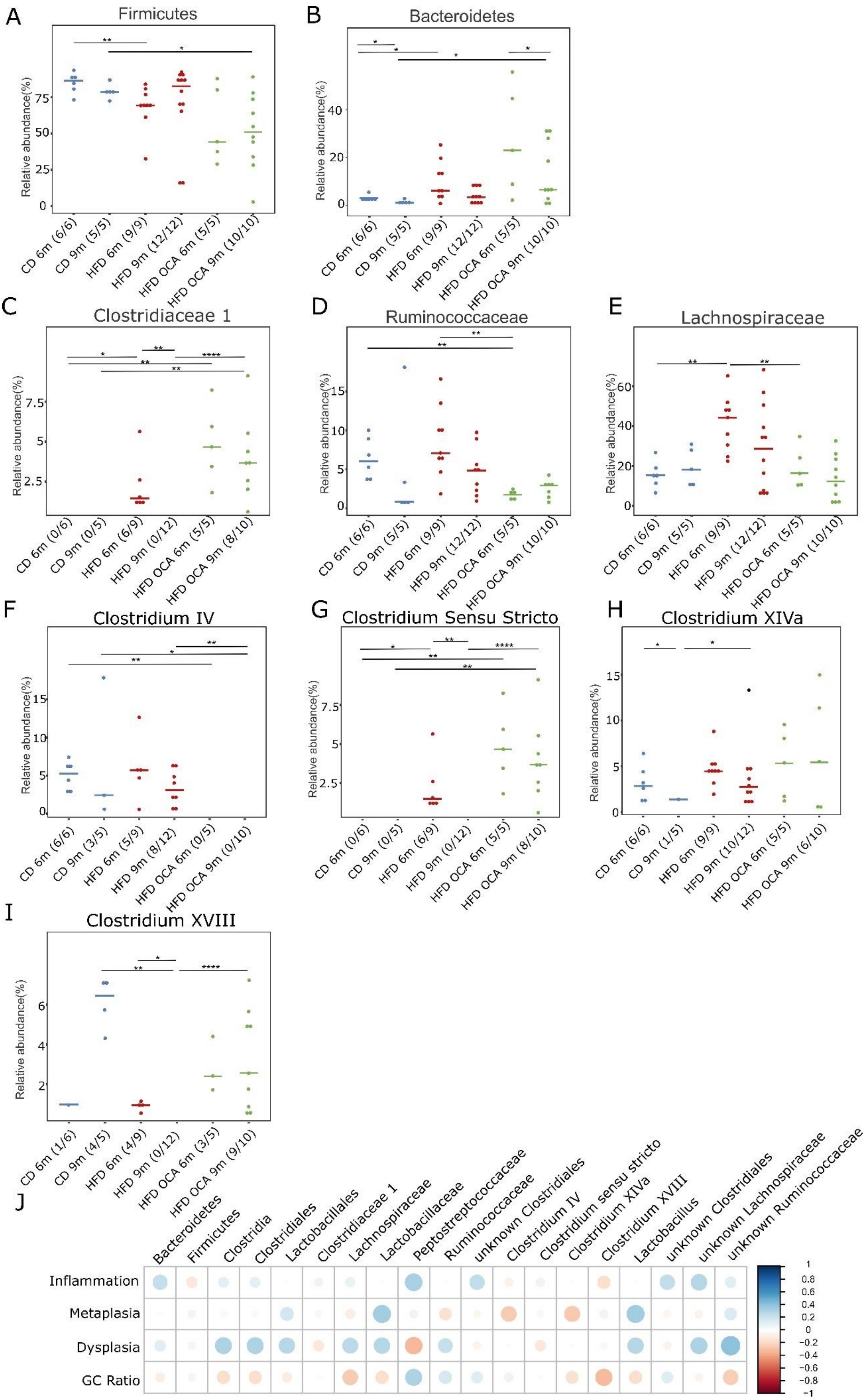

A
Firmicutes
Relative abundance(%)
CD 6m (6/6)
CD 9m (5/5)
HFD 6m (9/9)
HFD 9m (12/12)
HFD OCA 6m (5/5)
HFD OCA 9m (10/10)
B
Bacteroidetes
Relative abundance(%)
CD 6m (6/6)
CD 9m (5/5)
HFD 6m (9/9)
HFD 9m (12/12)
HFD OCA 6m (5/5)
HFD OCA 9m (10/10)
C
Clostridiaceae 1
Relative abundance(%)
CD 6m (0/6)
CD 9m (0/5)
HFD 6m (6/9)
HFD 9m (0/12)
HFD OCA 6m (5/5)
HFD OCA 9m (8/10)
D
Ruminococcaceae
Relative abundance(%)
CD 6m (6/6)
CD 9m (5/5)
HFD 6m (9/9)
HFD 9m (12/12)
HFD OCA 6m (5/5)
HFD OCA 9m (10/10)
E
Lachnospiraceae
Relative abundance(%)
CD 6m (6/6)
CD 9m (5/5)
HFD 6m (9/9)
HFD 9m (12/12)
HFD OCA 6m (5/5)
HFD OCA 9m (10/10)
F
Clostridium IV
Relative abundance(%)
CD 6m (6/6)
CD 9m (3/5)
HFD 6m (5/9)
HFD 9m (8/12)
HFD OCA 6m (0/5)
HFD OCA 9m (0/10)
G
Clostridium Sensu Stricto
Relative abundance(%)
CD 6m (0/6)
CD 9m (0/5)
HFD 6m (6/9)
HFD 9m (0/12)
HFD OCA 6m (5/5)
HFD OCA 9m (8/10)
H
Clostridium XIVa
Relative abundance(%)
CD 6m (6/6)
CD 9m (1/5)
HFD 6m (9/9)
HFD 9m (10/12)
HFD OCA 6m (5/5)
HFD OCA 9m (6/10)
I
Clostridium XVIII
Relative abundance(%)
CD 6m (1/6)
CD 9m (4/5)
HFD 6m (4/9)
HFD 9m (0/12)
HFD OCA 6m (3/5)
HFD OCA 9m (9/10)
J
Bacteroidetes
Firmicutes
Clostridia
Clostridiales
Lactobacillales
Clostridiaceae 1
Lachnospiraceae
Lactobacillaceae
Peptostreptococcaceae
Ruminococcaceae
unknown Clostridiales
Clostridium IV
Clostridium sensu stricto
Clostridium XIVa
Clostridium XVIII
Lactobacillus
unknown Clostridiales
unknown Lachnospiraceae
unknown Ruminococcaceae
Inflammation
Metaplasia
Dysplasia
GC Ratio

**Supplementary Figure 6: Representation of the cecal relative abundance (%) of bacterial families with BA metabolizing capacities between L2-IL1B mice fed with CD, HFD or HFD+OCA at 6 month and 9 month of age.**

Only relevant significant changes are shown.

- **(A)** The phylum Firmicutes decreased in abundance in 6-m-old HFD- compared to CD-fed mice and in 9-m-old HFD+OCA- compared to CD-fed mice (CD 6m vs HFD 6m p=0.0076; CD9 m vs HFD+OCA 9m p=0.0400).
- **(B)** The phylum Bacteroides was significantly higher abundant in 6-month-old HFD- and HFD+OCA-fed compared to CD-fed mice. The relative abundance droped between 6-m- and 9-m- in CD- and HFD+OCA-fed mice (CD 6m vs CD9 m p=0.0303; CD 6m vs HFD 6m p=0.0256; CD 9m vs HFD+OCA 9m p=0.0400; HFD+OCA 6m vs HFD+OCA 9m p=0.0400).
- **(C)** The family *Clostridiaceae 1* were only abundant in 6-m-old HFD-fed and 6-m- and 9-m-old HFD+OCA-fed mice (CD 6m vs HFD 6m p=0.0278; CD 6m vs HFD+OCA 6m p=0.0022; CD 9m vs HFD+OCA 9m p=0.0070; HFD 9m vs HFD+OCA 9m p=0.0001; HFD 6m vs HFD 9m p=0.0015).
- **(D)** The family of *Ruminococcaceae* was significantly decreased in abundance in 6m old HFD+OCA-fed mice compared to CD- and HFD-fed mice (CD 6m vs HFD+OCA 6m p=0.0043; HFD 6m vs HFD+OCA 6m p=0.0040).
- **(E)** The family of *Lachnospiraceae* was significantly enriched in abundance in 6-m-old HFD-fed mice compared to CD- and HFD+OCA-fed mice (CD 6m vs HFD 6m p=0.0016; CD 9m vs HFD 6m p=0.0120).
- **(F)** The genus *Clostridium cluster IV* was only abundant in CD- and HFD-fed mice (CD 6m vs HFD+OCA 6m p=0.0022; CD 9m vs HFD+OCA 9m p=0.0220; HFD 9m vs HFD+OCA 9m p=0.0017).
- **(G)** The genus *Clostridium sensu stricto* was only abundant in 6-m-old HFD-fed and in 6-m- and 9-m-old HFD+OCA-fed mice (CD 6m vs HFD 6m p=0.0278; CD 6m vs HFD+OCA 6m p=0.0022; CD 9m vs HFD+OCA 9m p=0.0070; HFD 9m vs HFD+OCA 9m p=0.0001; HFD 6m vs. HFD 9m p=0.0015).
- **(H)** Abundance of the genus *Clostridium cluster XIVa* was increased in 9-m-old HFD- vs CD-fed mice (p=0.0276) and decreased in 9-m- vs. 6-m-old CD-fed mice (p=0.0152).
- **(I)** The genus *Clostridium cluster XVIII* was significantly higher abundant in 9-m-old CD- and HFD+OCA-fed compared to HFD-fed mice (CD 9m vs HFD 9m p=0.0021; HFD 9m vs HFD+OCA 9m p<0.0001) and was not present in 9-m-old HFD-fed mice (HFD 6m vs HFD 9m p=0.0211).

**(J)** Correlation analysis between the abundance of bacterial families with BA metabolizing capacities in all treatment groups vs. inflammation, metaplasia, and dysplasia scores as well as goblet cell (GC) ratio. Correlation analyses were performed using the Rhea pipeline for 16s-sequencing data. *Clostridia, Lactobacillales, Ruminococcaceae* and unknown *Lachnospiraceae* were the main microbial groups positively correlated with dysplasia scores.

For (A-J) relative abundances of microbiota are represented as mean. For statistical analysis, pairwise Wilcoxon rank sum test or Fisher´s exact test, if appropriate, were used.

A

stool

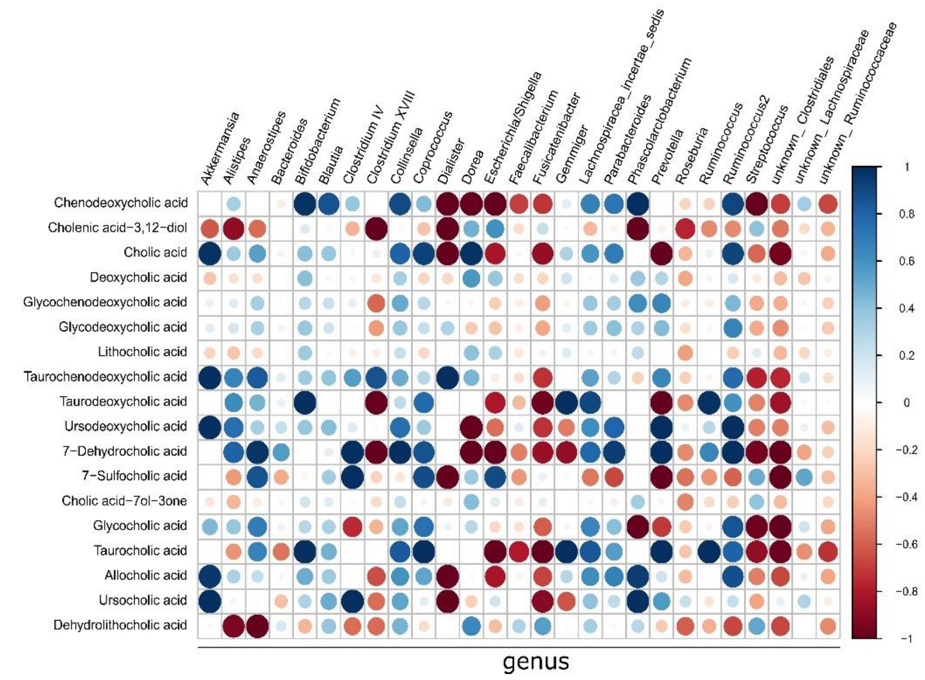


**Supplementary Figure 7: Intestinal microbial profile at genus level in relation to BA levels in patients**

**(A)** Correlation of the relative abundance of bacteria genera in the stool of human patients with disease severity and fecal BA levels. Correlation analyses were performed using the Rhea pipeline for 16S rRNA sequencing data.
